# CellTools algorithm for mapping scRNA-seq query cells to the reference dataset improves the classification of resilient and susceptible retinal ganglion cell types

**DOI:** 10.1101/2021.10.15.464552

**Authors:** Bruce A. Rheaume, Jian Xing, William C. Theune, Samantha G. DeRosa, Ephraim F. Trakhtenberg

**Affiliations:** Department of Neuroscience, University of Connecticut School of Medicine, 263 Farmington Ave., Farmington, CT, 06030, USA

## Abstract

The clustering of single cell RNA-sequencing (scRNA-seq) data enables the classification of cell types, and the development of integration mapping algorithms has enabled the tracing of altered-from-baseline transcriptomes to their respective cell type origins. Here, we developed an algorithm that removes sources of noise from scRNA-seq reference dataset, and in the next step optimizes weight-assignment to anchors, cumulatively improving the accuracy of query cells mapping to reference dataset. The denoising step of our algorithm also improved the performance of other mapping algorithms. To further demonstrate biological relevance, using our algorithm we determined the type-origin of the 17% of injured retinal ganglion cells (RGCs) that a prior algorithm did not identify. As we found that most of the originally unassigned cells belonged to only some RGC types, a consequent change in the proportions of the surviving types resulted in an amended ranking of resiliency to injury. We also identified new cluster-markers for RGC types, validated two novel markers by immunostaining in retinas, and developed a website for cluster-by-cluster comparison of gene expression between uninjured and injured RGC types. Additional bioinformatic analyses contributed new insights into the global characteristics of RGC types and how axonal injury affects them, showing how dissimilarity between transcriptomes of RGC types increases during maturation and after injury. We further characterized the correspondence between the neonatal and adult RGC types, and showed which cluster markers change expression developmentally or after injury. We also show, for the first time, that global properties of the transcriptome can predict the resilience to injury of at least some cell types. The R-package, CellTools, for the algorithms we developed, will assist scRNA-seq studies across biological fields.

## INTRODUCTION

The existing integration mapping algorithms, widely used for tracing altered transcriptomes of matured and/or injured cells to their respective scRNA-seq cluster-defined cell type origin, are evolving as new engineering and computational approaches emerge in the field. However, the limitations of comprehensiveness and accuracy of cell-to-cluster assignment by the existing integration mapping algorithms requires new advancements^1-3^. Here, we developed CellTools algorithms, which first denoise the scRNA-seq reference dataset, and then determine the optimal parameters for reference-driven mapping integration of query scRNA-seq dataset. Our approach improved quality of the reference dataset (by identifying poor quality cells) and increased the percent of correct ‘query cell’-to-‘reference cluster’ assignments, as compared to several other scRNA-seq integration mapping algorithms.

For a proof-of-concept of that a more accurate integration mapping algorithm can effect biological conclusions, we show that implementation of our algorithm improved the identification of scRNA-seq-derived resilient and susceptible retinal ganglion cell (RGC) types, which were originally classified using the integration mapping algorithm iGraphBoost^4^. We also introduced an algorithm for analyzing global properties of the transcriptome and show for the first time that it can predict biological phenotypes, such as the resiliency to injury of at least some RGC types. Furthermore, we identified new gene cluster markers for RGC types, contributed new insights into how dissimilarity between transcriptomes of RGC types increases during maturation and after injury, and developed a website for cluster-by-cluster comparison of gene expression between uninjured and injured RGC types. We made the CellTools R software library for our algorithms publicly available.

## RESULTS

### Retinal ganglion cell scRNA-seq datasets used for CellTools algorithms development

Mouse RGCs are comprised of 40-46 types^4-8^ (sometimes referred to as subtypes^5,9-11^), with some types surviving longer after optic nerve crush (ONC) injury and/or being more capable of regenerating axons^4,12,13^. Classification of the uninjured adult mouse RGC scRNA-seq-derived transcriptome profiles (adult RGC atlas, which we used for developing our algorithm) by the Louvain-Jaccard algorithm^14,15^ predicted 45 clusters, one of which was manually sub-clustered into 2, resulting in a total of 46 types^4^, that closely matched the 40 neonatal RGC types we identified previously^5^ (see Supplementary Fig. 1F in Tran et al., 2019^4^). However, the iGraphBoost algorithm traced the cluster origin of only 82.8% of the injured RGCs (that survived 2 weeks after ONC, which is a standard time-point in this assay^4,16^) to the uninjured RGC types, despite the data from earlier time-points (i.e., before 2 weeks after ONC) enabling sequential tracing of RGC types, as their transcriptomes were fluctuating over time after injury (see Fig. 3C in Tran et al., 2019^4^). The resilience or susceptibility of RGC types was, in turn, determined by the prevalence of the types remaining 2 weeks after injury compared to the prevalence of their uninjured counterpart types (see Fig. 3E in Tran et al., 2019^4^). The implicit assumption of such an analysis is that specific RGC types were not overrepresented amongst the unaccounted 17.2% of injured RGCs; otherwise, such types would have been deemed susceptible when in fact they are resilient. Because accurate determination of the resilient and susceptible RGC types depends on this assumption, we selected this dataset for validating our algorithm by testing whether it could improve cell-to-cluster assignment in the integration mapping and thereby enable testing this assumption.

### Two-step algorithm for improving accuracy of mapping integration

To improve query-to-reference scRNA-seq datasets integration mapping, we developed a two-step algorithm, which first denoises reference dataset, and then optimizes the *k*-weight parameter of the K-nearest neighbor (KNN) algorithm for a more accurate cell-to-cluster assignment (using RGC scRNA-seq dataset, see above and in detail below; **Fig. 1A-K** and **Supplemental Figure 1A-C**). We show significant improvement by each step, using subsampling of the atlas of adult uninjured RGC types as a proof-of-concept (**Fig. 1M**), which cumulatively reduced incorrect integration (i.e., mapping to reference) of query cells by 73% (on average).

**Figure 1.**
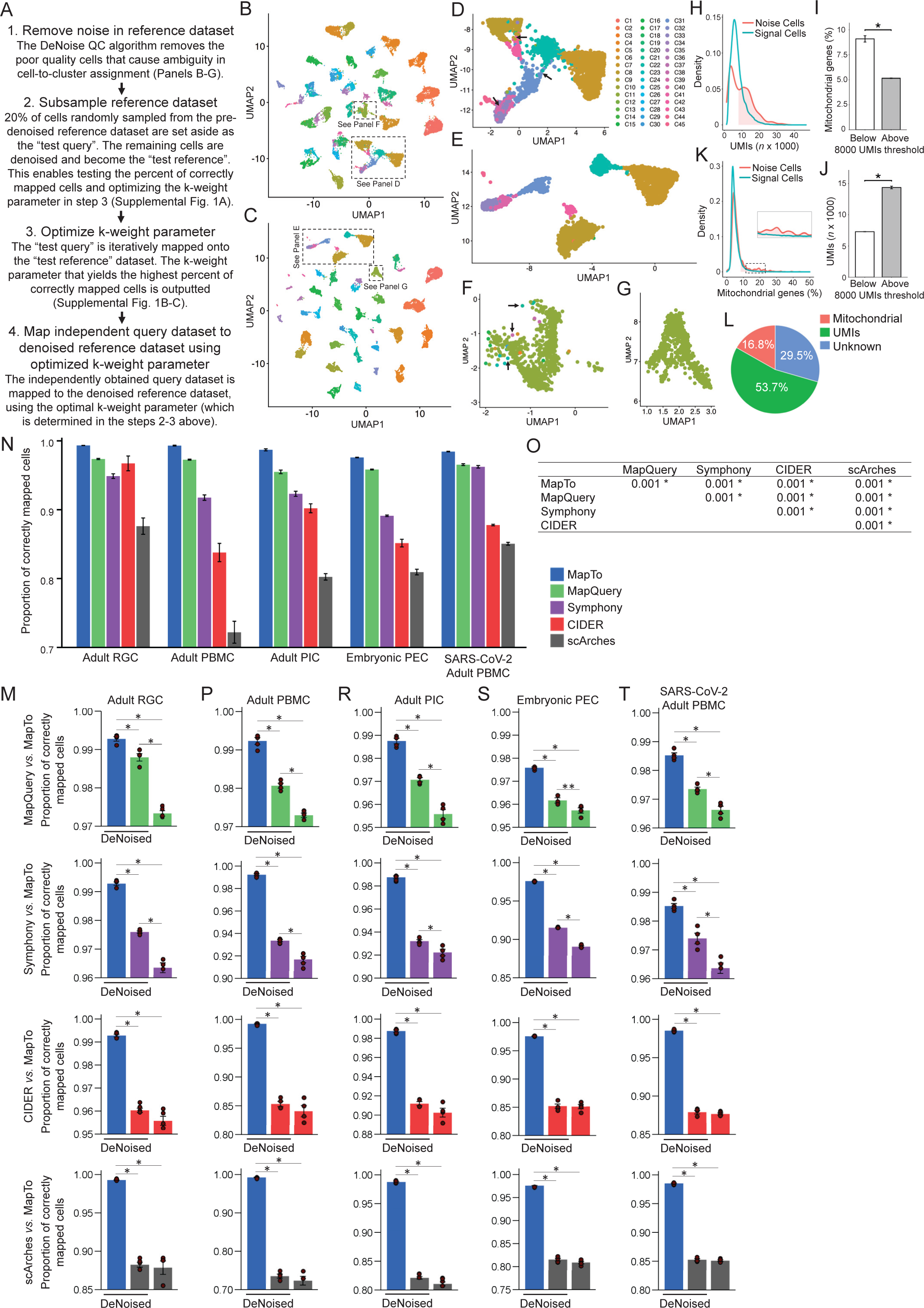
CellTools algorithms remove sources of noise and improve accuracy of mapping integration. (**A**) A flowchart of the steps that the two-step algorithm performs. First, the noise cells are removed from the reference dataset by the DeNoise function. Next, 20% of the randomly subsampled reference dataset is set aside as a “test query” and the remainder becomes the “test reference”. Then, the test query is iteratively mapped over a range of *k*-weights (see methods) to the test reference, and the *k*-weight that yields the highest percent of correctly mapped cells is selected as optimal for mapping the original query dataset. (**B-C**) The UMAP of uninjured adult atlas RGCs (***B***) yielded a similar distribution of the clusters as the tSNE (colored by Louvain Jaccard-determined clusters) that was used originally (see Fig. 1C in Tran et al., 2019^6^). Clusters in all UMAPs and bar plots (*B-G, N-O,* and *R-S*) have the same color-code, shown in the legend to the right of (*D*). A denoised reference dataset UMAP (***C***) shows improved cluster-UMAP agreement (compared to B). (**D-G**) Insets (outlined in *B*-*C*) show reduction in inter-cluster heterogeneity (***D*** compared to improved ***E***), and increase in intra-cluster homogeneity (***F*** compared to improved ***G***). Arrows point to examples of ambiguity that are resolved by the DeNoise algorithm. (**H-K**) A unimodal distribution in density of the UMIs of the “signal” (i.e., not noise) cells overlapped the first peak in the bimodal distribution of the “noise” cells (***H***), but the second peak representing an abnormally large transcriptome size was unique to the “noise” cells (***I***). However, the first (normal) peak of UMIs of the “noise” cells (in *H*, annotated “below” in *I*) was significantly enriched for mitochondria-related genes (***J***). Conversely, a second peak representing abnormally high proportion of mitochondria-related genes was also unique to “noise” cells on the mitochondria-related genes density plot, whereas “signal” cells unimodal distribution overlapped the first peak in the bimodal distribution of the “noise” cells (***K***). * *p* < 0.0001 by *t*-test, 2-tailed; error bars = SEM. (**L**) A color-coded pie graph of the “noise” cells, showing abnormalities in 83.2% of these cells accounted for by abnormally large transcriptome size (53.7%) or abnormally high proportion of mitochondria-related genes (29.5%). (**M**) Average (from using different groups of randomly subsampled set-aside cells, *N* = 4) percent of “test query” RGC cells correctly mapped to the “test reference” RGC dataset by default integration (MapQuery) is significantly improved by denoising the reference dataset (DeNoised+MapQuery), and further improved by optimizing the *k*-weight parameter (DeNoised+MapTo), as shown in the upper panel. Lower panels show comparisons of our 2-step algorithm (DeNoise followed by MapTo) and the other integration mapping algorithms with or without application of the DeNoise algorithm, as marked. Significant *p*-values are indicated by an asterisk (*); * *p* <0.01 by ANOVA with posthoc LSD; error bars = SEM. (**N-O**) Comparative analysis of the average (from using different groups of randomly subsampled set-aside cells, *N* ≥ 3) percent of “test query” cells correctly mapped to the “test reference” dataset, for each of the five cell types’ scRNA-seq datasets by different integration mapping algorithms, as marked (***N***). The difference between the algorithms was significant by ANOVA (overall *F* = 695.4, *p* < 0.001), and pairwise comparisons by posthoc LSD showed significantly higher (*p* < 0.001, main effects) proportion of correctly mapped cells by our 2-step algorithm (DeNoise followed by MapTo), compared to each of the other integration mapping algorithms on multiple cell types datasets, as marked (***O***). Significant *p*-values are indicated by an asterisk (*); error bars = SEM. Cell types: RGC = Retinal ganglion cell; PBMC = Peripheral blood mononuclear cell; PIC = Pancreatic islets cell; PEC = Pancreatic epithelial cell; SARS-CoV-2 PBMC = SARS-CoV-2-infected PBMC. (**P-T**) Comparative analysis of the average (from using different groups of randomly subsampled set-aside cells, *N* ≥ 3) percent of “test query” cells correctly mapped to the “test reference” dataset, for each of the five cell types’ scRNA-seq datasets by different integration mapping algorithms, with or without application of the DeNoise algorithm first, as marked. The application of the DeNoise algorithm improved significantly (*p* < 0.02 by ANOVA with posthoc LSD) the performance of Seurat’s MapQuery and Symphony integration mapping algorithms, compared to their performance without the application of the DeNoise algorithm, on all five cell types datasets. Nevertheless, our 2-step algorithm (DeNoise followed by MapTo) performed significantly (*p* < 0.01 by ANOVA with posthoc LSD) better, compared to each of the other integration mapping algorithms on all five cell types datasets, even if they were applied after the application of our DeNoise algorithm first. Significant *p*-values are indicated by an asterisk (*): * *p* <0.01, ** *p* <0.02; error bars = SEM.

Our algorithm applies a wrapper for the Seurat functions (FindTransferAnchors, TransferData, IntegrateEmbeddings, and ProjectUMAP)^1,17^, which use high-dimensional data for determining the integration anchors used for mapping the query cells, and also uses uniform manifold approximation and projection (UMAP)^18,19^ for transferring the reference cluster labels of nearest neighbors, that are determined, in part, by the *k*-weight parameter of the KNN algorithm. UMAP yielded a similar distribution of the Louvain Jaccard-determined clusters (**Fig. 1B**) as did the scalable implementation of t-distributed stochastic neighbor embedding (tSNE)^20,21^ that was used originally (see Fig. 1C in Tran et al., 2019^4^).

In the first step, the DeNoise function we developed, utilizes post-clustering structure of the dataset to identify and remove poor quality cells from the reference dataset, and then generates a denoised reference UMAP (**Fig. 1C**; see Methods) for mapping integration of the query cells in the next step. The DeNoise function reduces ambiguity in cluster segregation (inter-cluster heterogeneity; **Fig. 1D-E**) and increases intra-cluster homogeneity in the reference dataset (**Fig. 1F-G**), both of which impact weight-assignment of the anchors used for mapping integration of the query cells to the reference dataset clusters.

The DeNoise algorithm identified ∼3% of the total reference dataset cells as noise, and its validity was demonstrated by subsequent analyses, which revealed abnormalities in 83.2% of the noise cells (**Fig. 1H-L**). For example, 53.7% of the noise cells had an abnormally large transcriptome size (**Fig. 1H-J**), which could result from a partially damaged or a smaller cell binding to the same bead as another intact cell (and pass the doublet QC because the damaged or a smaller cell did not preserve its entire transcriptome), and 29.5% of the other noise cells had an abnormally high proportion of mitochondria-related genes (**Fig. 1I-K**), which is typically higher in cells progressing towards apoptosis^1,22,23^ (these cells could have been damaged more than other cells during processing). However, noise cells were not overrepresented in any clusters (**Supplemental Figure 2A**), and the clusters before and after denoising were overall highly similar (*r* = 0.99, Pearson); mean correlation coefficients of the pre-denoise and denoised clusters compared to every other atlas RGC cluster (pre-denoise and denoised, respectively) were also highly similar (average *r* = 0.99, Pearson; **Supplemental Figure 2B**). Our approach of identifying outliers based on the post-clustering structure of the dataset enabled removing sources of noise not detected by a doublet-removal algorithm (Scrublet^24^), which predicted only one doublet in the pre-denoised reference dataset (which is not surprising as we used published atlas RGC cells that already passed QCs). Moreover, the predicted doublet was not amongst the “noise” cells detected by our algorithm, as it did not present as an outlier in the post-clustering structure of the dataset and its UMI was within 1 SD of the respective cluster mean UMI (**Supplemental Figure 2C-D**). Further validation of the DeNoise algorithm was shown by that it has significantly improved the integration mapping of the query cells, by merely eliminating sources of noise from the reference dataset, even before machine learning algorithm optimization of the *k*-weight parameter in the next step (**Fig. 1M** *upper panel*).

In the second step, the MapTo function improves the accuracy of cell-to-cluster assignment by choosing the *k*-weight parameter of the KNN algorithm in a data-driven manner. The *k*-weight parameter is the number of neighbors (i.e., cells) the KNN algorithm considers when assigning weights to anchors (i.e., neighboring pairs of cells from the reference and query datasets) in determining cell-to-cluster assignment. Our machine learning algorithm optimizes the *k*-weight parameter for the reference dataset, by randomly setting aside 20% (a default that can be changed by user) of the total cells from the reference dataset before clustering it, and determining at which *k*-weight parameter the maximum percent of these cells are accurately traced by the integration mapping algorithm to the type-origin clusters (i.e., clusters to which the set-aside cells belonged originally) (**Supplemental Figure 1A**). To determine the optimal *k*-weight parameter, which yields the highest percent of accurately traced set-aside cells, the algorithm uses a loop-script (see Methods) that iterates through the Seurat functions’ *k*-weight parameters and outputs the percent of the accurately traced set-aside cells. The algorithm then ranks the percent of the accurately traced cells from all iterations, and selects the *k*-weight parameter which yields the highest percent of correctly-mapped cells (**Supplemental Figure 1B-C**). We repeated this process 3 times, using different groups of randomly subsampled set-aside cells, and found that the optimal *k*-weight parameter significantly improved the accuracy of their mapping to the reference (**Fig. 1M** *upper panel*).

### Comparative analysis on the accuracy of mapping integration using other algorithms and datasets

To further validate our 2-step algorithm (DeNoise followed by MapTo), we comparatively evaluated its performance to more integration mapping algorithms (CIDER^25^, scArches^26^, and Symphony^27^, in addition to Seurat’s MapQuery^17^) and on five different scRNA-seq datasets: Adult mouse retinal ganglion cells (RGC) used herein in the initial validation of the 2-step (MapTo and DeNoise), adult human peripheral blood mononuclear cells (PBMC) used in validation of the Seurat’s MapQuery^17^, adult human SARS-CoV-2-infected PBMCs used in validation of the Symphony^27^, adult mouse and human pancreatic islets cells (PIC) used in validation of CIDER^25^, and embryonic mouse pancreatic epithelial cell (PEC) used in validation of scArches^26^. As above, we performed the analysis at least 3 times on each dataset, using different groups of randomly subsampled set-aside cells each time (in order to account for technical variability). We found that significantly (*p* < 0.001) higher accuracy of mapping to the reference was achieved by our 2-step algorithm, as compared to the other integration mapping algorithms, on each of the five different cell types datasets (**Fig. 1N-O** and **Table 1**). We also found that Seurat’s MapQuery^17^ and Symphony^27^ integration mapping algorithms performed significantly (*p* < 0.02) better, on each of the five different cell types datasets, if applied after the DeNoise module (which is the first step of our algorithm), suggesting that the DeNoise algorithm on its own can be compatible with other mapping algorithms (**Fig. 1M,P-T**).

**Table 1.** Pairwise comparisons for the interaction effects of mapping algorithms and cell types. Pairwise comparisons by posthoc LSD for the interaction effects of mapping algorithms and cell types from the Fig. 1N ANOVA analysis, showing *p*-values for each comparison. The difference between the interaction effect of algorithms and cell types was significant by ANOVA (overall *F* = 27.4, *p* < 0.001), and pairwise comparisons by posthoc LSD showed significantly higher (*p* < 0.05, interaction effects) proportion of correctly mapped cells by our 2-step algorithm (DeNoise followed by MapTo), compared to each of the other integration mapping algorithms on every cell type dataset, as marked. Significant *p*-values are indicated by an asterisk (*). RGC = Retinal ganglion cell; PBMC = Peripheral blood mononuclear cell; PIC = Pancreatic islets cell; PEC = Pancreatic epithelial cell; SARS-CoV-2 PBMC = SARS-CoV-2-infected PBMC.

| Adult RGC |  |  |  |  |
| --- | --- | --- | --- | --- |
|  | MapQuery | Symphony | CIDER | scArches |
| MapTo | 0.022 * | 0.0001 * | 0.003 * | 0.0001 * |
| MapQuery |  | 0.004 * | 0.447 | 0.0001 * |
| Symphony |  |  | 0.031 * | 0.0001 * |
| CIDER |  |  |  | 0.0001 * |

| Adult PBMC |  |  |  |  |
| --- | --- | --- | --- | --- |
|  | MapQuery | Symphony | CIDER | scArches |
| MapTo | 0.018 * | 0.0001 * | 0.0001 * | 0.0001 * |
| MapQuery |  | 0.0001 * | 0.0001 * | 0.0001 * |
| Symphony |  |  | 0.0001 * | 0.0001 * |
| CIDER |  |  |  | 0.0001 * |

| Adult PIC |  |  |  |  |
| --- | --- | --- | --- | --- |
|  | MapQuery | Symphony | CIDER | scArches |
| MapTo | 0.0001 * | 0.0001 * | 0.0001 * | 0.0001 * |
| MapQuery |  | 0.0001 * | 0.0001 * | 0.0001 * |
| Symphony |  |  | 0.015 * | 0.0001 * |
| CIDER |  |  |  | 0.0001 * |

| Embryonic PEC |  |  |  |  |
| --- | --- | --- | --- | --- |
|  | MapQuery | Symphony | CIDER | scArches |
| MapTo | 0.041 * | 0.0001 * | 0.0001 * | 0.0001 * |
| MapQuery |  | 0.0001 * | 0.0001 * | 0.0001 * |
| Symphony |  |  | 0.0001 * | 0.0001 * |
| CIDER |  |  |  | 0.0001 * |

Table 1. Pairwise comparisons for the interaction effects of mapping algorithms and cell types
| Adult SARS-CoV-2 PBMC |  |  |  |  |
| --- | --- | --- | --- | --- |
|  | MapQuery | Symphony | CIDER | scArches |
| MapTo | 0.027 * | 0.011 * | 0.0001 * | 0.0001 * |
| MapQuery |  | 0.711 | 0.0001 * | 0.0001 * |
| Symphony |  |  | 0.0001 * | 0.0001 * |
| CIDER |  |  |  | 0.002 * |

### Assignment of previously unsigned injured RGCs to atlas RGC types

We then re-analyzed the original data using the algorithm we developed, in order to test whether 17.2% of the injured RGCs were not traced to respective cluster origin due to the computational bioinformatic limitations, or due to biological changes after injury that dedifferentiated certain RGCs beyond bioinformatic traceability to types. We found that our algorithm enabled the tracing of all the RGCs to their type-origin (**Fig. 2A-B**; at 2 weeks after ONC), and increased the correlation between the uninjured and the injured RGC clusters (traced to respective type-origin) for the analyzed post-injury time-points, compared to using the original analyses (**Supplemental Figure 3A**). For the 2 weeks after ONC time-point types assigned by the original analysis and by our algorithm were overall highly similar (average *r* = 0.98, Person), despite the inclusion of the additional 17.2% of the originally unassigned RGCs now assigned to types by our algorithm. We also found that, injured RGC clusters (after assignment; **Supplemental Figure 3A**), as well as all individual injured RGCs (pre-assignment; **Supplemental Figure 3B**), correlated with the uninjured RGC clusters or all individual RGCs, respectively, non-linearly relative to the analyzed post-injury time-points. For example, RGCs at 7 days after ONC were slightly (but significantly, *p* < 0.001) more similar to uninjured RGCs than the RGCs from a preceding time-point at 4 days after ONC, despite progressive degeneration of the injured RGCs over time (**Supplemental Figure 3A-B**). There was also a peak in the percentage of injured RGCs unassigned to clusters by iGraphBoost at 4 and 7 days after ONC, compared to a smaller percent of RGCs unassigned at earlier time-points (at 1 and 2 days after ONC) and a later time-point (at 2 weeks after ONC; **Supplemental Figure 3C**), suggesting increased fluctuations in some RGCs’ gene expression profiles at different time-points after injury that rendered more cells unassigned at earlier compared to later time-points in the original analysis (which is also consistent with that the highest variably between the batches was at 4 days after ONC; **Supplemental Figure 3D**). Therefore, using the sequential (i.e., iterative integration) tracing may not be advantageous for the 2-week post-ONC time-point *per se*, but the time-points dataset is important for determining early and late susceptibility types (**Supplemental Figure 3E-H**).

**Figure 2.**
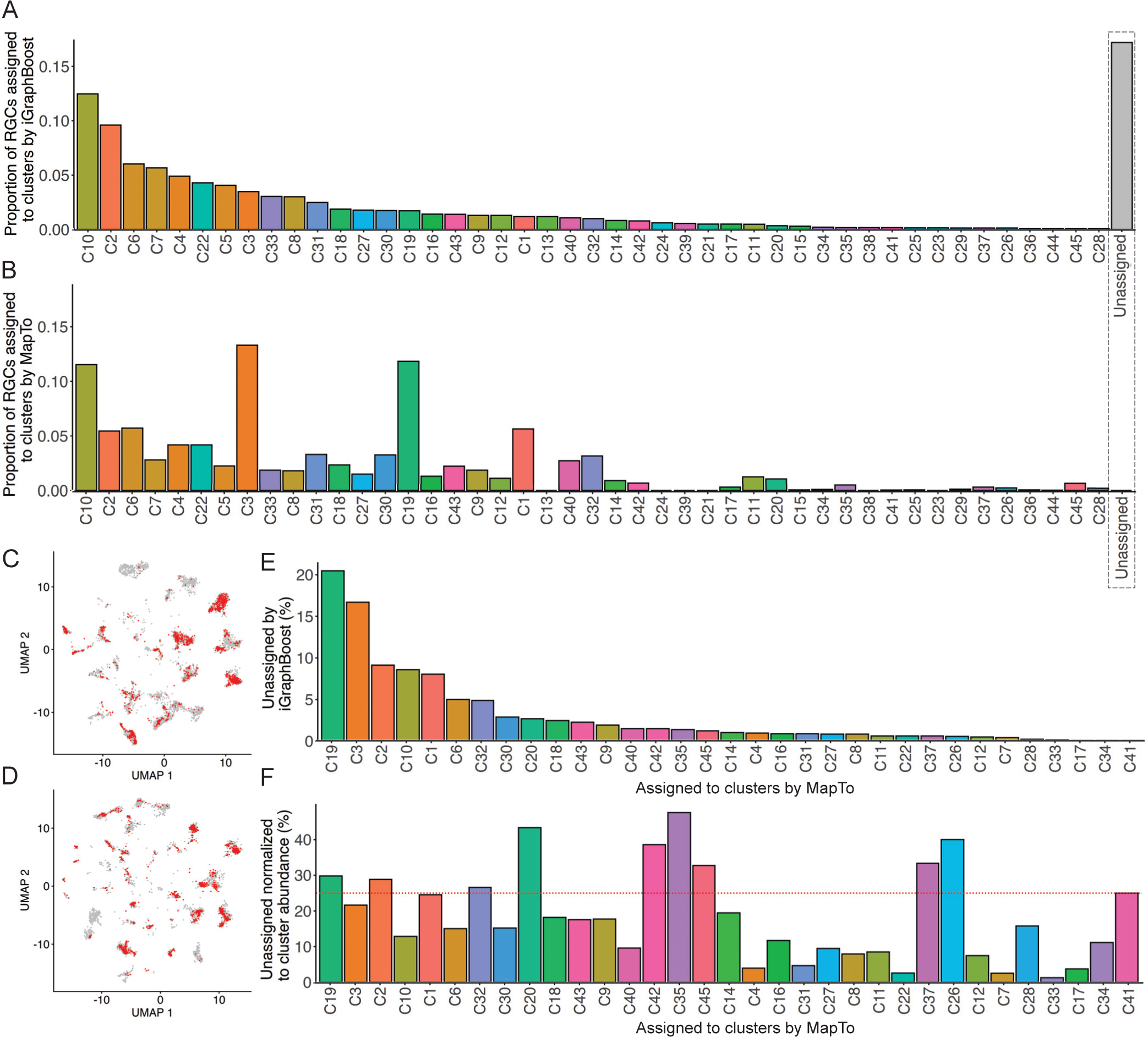
Overrepresentation of originally unassigned injured RGCs in a subset of clusters. (**A-B**) Proportion of RGCs at 2 weeks after ONC assigned to clusters by the iGraphBoost (***A***) and two-step MapTo (***B***) algorithms, showing a substantial percent of the injured RGCs unassigned to clusters by the iGraphBoost algorithm and the absence of unassigned RGCs by the two-step MapTo algorithm (bar representing the unassigned RGCs outlined by dashed line). Clusters in *A* and *B* are ordered (left to right) following higher to lower proportion in *A*. (**C-D**) UMAPs of injured RGCs (at 2 weeks after ONC) assigned by the two-step MapTo algorithm to uninjured atlas reference dataset, before (***C***; mapped to the original reference dataset shown in Fig. 1B) and after denoising (***D***; mapped to the denoised reference dataset shown in Fig. 1C). Color-coded (in Fig. 1C-D) previously assigned (gray) and unassigned (red) by the iGraphBoost algorithm cells, help to visualize that originally unassigned injured RGCs (red cells) remained overrepresented in certain clusters even after denoising (*D*). (**E-F**) The two-step MapTo algorithm assignments of RGCs previously unassigned by the iGraphBoost algorithm (at 2 weeks after ONC) as percent of total originally unassigned RGCs (***E***), and as percent of total RGCs assigned per respective cluster (***F***), showing substantial (>25%) overrepresentation in clusters C2, C19, C20, C26, C32, C35, C37, C42, and C45. Clusters in *E* and *F* are ordered (left to right) following higher to lower percent in *E*.

Next, we analyzed whether certain RGC types were overrepresented amongst the originally unassigned injured RGCs. We found that the majority (63%) of the originally unassigned RGCs belonged to only five types (C1, C2, C3, C10, and C19), and that types C2, C19, C20, C26, C32, C35, C37, C42, and C45 were substantially (>25% each) overrepresented amongst the originally unassigned RGCs (**Fig. 2C-F**). Then, we analyzed the prevalence of the surviving RGC types, with all injured RGCs now assigned to types, and found that determination of several resilient and susceptible RGC types changed compared to their original ranking (see Fig. 3E in Tran et al., 2019^4^). For example, the originally susceptible types C45, C19, and C3 now rank as resilient, the originally susceptible type, C1, now ranks as neutral, and the originally neutral type, C39, now ranks as susceptible (**Fig. 3A-C**). Because αRGCs, to which C45 type belongs, were previously shown to be more resilient to injury^12,13,28,29^, the change of C45 rank from susceptible to resilient further supports the improved accuracy of our algorithm. We also developed the Subtypes Gene Browser web application (in the same format we previously presented for neonatal RGC subtypes from the left and right eyes^5^), which provides a platform for cluster-by-cluster differential gene expression analysis between the atlas and injured RGC clusters (https://health.uconn.edu/neuroregeneration-lab/subtypes-gene-browser; see Methods for details).

**Figure 3.**
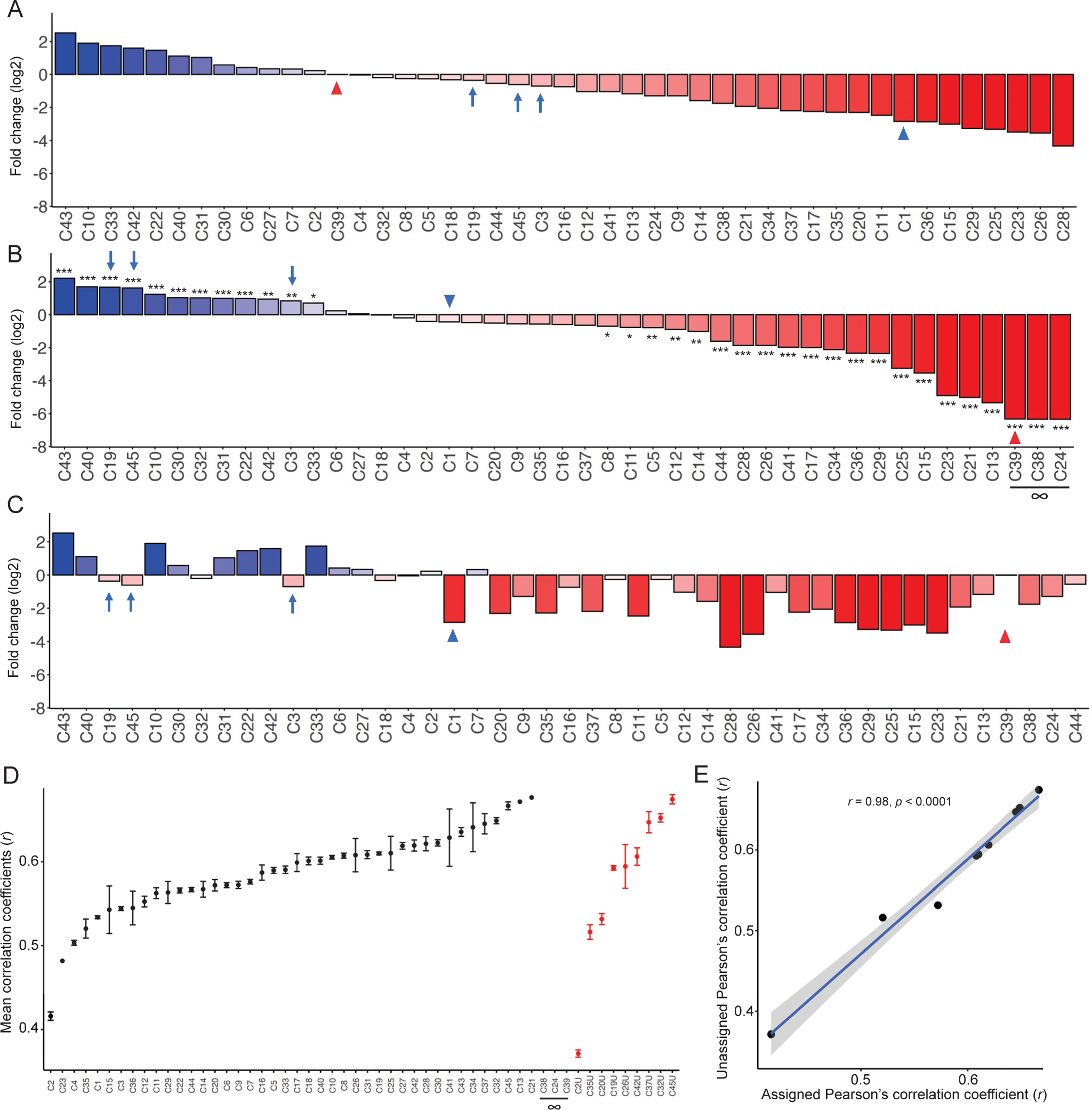
Re-analysis of the RGC types survivability after assignment to clusters of the previously unassigned injured RGCs. (**A-B**) RGC types’ resilience/susceptibility determined by log2 fold-change of uninjured atlas RGCs to injured RGCs (at 2 weeks after ONC) based on cell-to-cluster assignment using iGraphBoost (replicated from Fig. 3E in Tran et al., 2019^6^) (***A***) or two-step MapTo (***B***) algorithm. Fold change significance in (*B*) determined by EdgeR analysis, as indicated (\*\*\**p* < 0.001, \*\**p* < 0.01, and \**p* < 0.05; see Methods). (**C**) To better visualize the shifts in type resilience/susceptibility determination (i.e., the differences between *A* and *B*), the same cluster order as in *B* (MapTo-determined) is shown with the cluster log2 fold-change and color-code as in *A* (iGraphBoost-determined). Arrows and arrowheads point to the same clusters at different positions in the panels (*A*-*C*): Blue arrow and arrowhead indicate examples of clusters that changed determination from “susceptible” to “resilient” or to “neutral” (i.e., clusters that did not show significant change in *B*), respectively; red arrowhead indicates an example of a cluster that changed determination from “neutral” to “susceptible”. (**D**) Black datapoints: the mean correlation coefficients (*r*) of the injured (2 weeks after ONC) RGCs assigned to clusters by the two-step MapTo algorithm (excluding those that were originally unassigned to clusters by the iGraphBoost algorithm) and all uninjured atlas RGCs from their respective clusters. Red datapoints: Mean correlation coefficients (*r*) of the injured (2 weeks after ONC) RGCs that were originally unassigned to clusters by the iGraphBoost algorithm (but assigned to clusters in retrospect by the MapTo algorithm) and uninjured atlas RGCs from their respective clusters. C24, C38, and C39, which did not survive 2 weeks after injury are marked by infinity (∞). The range of distribution for the black and red datapoints is similar, indicating that the changes due to injury in the transcriptomes of originally unassigned RGC types are within the normal range compared to the other RGC types. U = Unassigned; Error bars = SEM. (**E**) Linear regression of mean correlation coefficients of the C2U, C19U, C20U, C26U, C32U, C35U, C37U, C42U, and C45U (that were originally unassigned to clusters by the iGraphBoost algorithm, but assigned to clusters in retrospect by the two-step MapTo algorithm) and uninjured atlas RGCs from their respective clusters (red datapoints in *D*), vs. mean correlation coefficients of injured RGCs assigned to counterpart clusters (excluding the RGCs in C2U, C19U, C20U, C26U, C32U, C35U, C37U, C42U, and C45U that were originally unassigned to clusters by the iGraphBoost algorithm) and uninjured atlas RGCs from their respective clusters (black datapoints in *D* for C2, C19, C20, C26, C32, C35, C37, C42, and C45). High correlation coefficient (*r* = 0.98; Pearson’s) indicates that the transcriptomes of originally unassigned RGC types are highly correlated with their assigned counterpart clusters. Gray area around the fit-line indicates 95% confidence interval for the linear regression model.

Then, we analyzed whether, at 2 weeks after ONC, the injured RGCs which were unassigned to types by the original algorithm (iGraphBoost) were substantially different from the injured RGCs that were assigned to types by our algorithm. We found that, the transcriptomes of the originally unassigned overrepresented injured RGC types, not only did not belong to the types most or least changed by injury (**Fig. 3D**), but when grouped separately based on our type-assignment were highly correlated (*r* = 0.98, *p* < 0.0001) to their originally assigned counterpart types (**Fig. 3E**), thereby further supporting the accuracy of their assignment to clusters by our algorithm. Moreover, expression pattern of the injured RGC type markers, which were enriched in the originally assigned RGC types, was similar in the types assigned by our algorithm to the originally unassigned RGCs (**Supplemental Figure 4**). We also did not find the relative proportion of mitochondria-related genes (see above) in the overrepresented unassigned RGC types to be amongst the highest or lowest relative to the assigned types (**Supplemental Figure 5A**), nor was the total number of genes detected in the originally unassigned injured RGC types amongst the highest or lowest relative to the assigned types (**Supplemental Figure 5B-C**). The overrepresented unassigned RGC types also were not amongst the least or most abundant types in the RGC atlas (**Supplemental Figure 5D**). These data suggest that the injured RGCs unassigned to types by the original algorithm (iGraphBoost) were not substantially different from other injured RGCs.

### Correspondence of neonatal and adult RGC types

Next, we characterized the correspondence between the classification of adult RGCs into 45 types^4^ and our prior classification of neonatal RGCs into 40 types^5^, after applying DeNoise to the neonatal RGC reference dataset, which identified ∼2.9% of cells as noise (52.5% of which were also flagged by a doublet-removal Scrublet^24^). Clustering together the neonatal and adult RGC datasets (see Methods) matched the different age types despite changes in gene expression associated with RGC maturation^30,31^ (**Fig. 4A-C**). While most neonatal and adult RGC types matched one-to-one, others diverged or converged into a few clusters (**Fig. 4D**). For example, the neonatal RGC type 16 split in adult into three Other (uncharacterized) types, 27, 37, and 44 (**Fig. 4E-F**). The neonatal Opn4+ ipRGC type 25 split in adult into three Opn4+ ipRGC types, 31, 33, and 40 (**Fig. 4G**). The remaining Opn4+ neonatal RGC types 6, 26, 37, and 33 (some of which were also independently validated^32^, and along with type 25 include the ipRGC types, together comprising intermediate subpopulation (ISP) 2^5^) matched the adult Opn4+ RGC types, 7, 8, 22, 31, 33, 40 (which include the ipRGC types) and 43 (which is an Opn4+ αRGC type). An Opn4-neonatal αRGC type 19 split in adult into two Opn4-RGC types, 41 αRGC and 18 Other (**Fig. 4H**). We also found that Tbr1-expressing (T-RGC) neonatal RGC types, 9, 14, 15, 21, and 38 (which comprised ISP 9^5^), matched adult T-RGC types 5, 9, 17, 21, and an αRGC type 42, which is also one of the five Tbr1-expersing types in adult RGCs. The remaining Opn4-/Tbr1-αRGC type 45 matched neonatal RGC type 39, which was also independently validated^32^. We also found that neonatal ooDSGC types 22 and 23 (which comprised ISP 1^5^) converged in adult into ooDSGC cluster 16 (**Fig. 4I**), possibly as a result of developmental pruning or circuitry refinement during maturation. The remaining adult ooDSGC types 10, 12, and 24 matched neonatal types 31, 32, and 29 (which were nested on different ISP branches 3-6 that stemmed from one of the four main branches of neonatal RGC subtypes diversification^5^). We also found that, neonatal Neurod2-expressing RGC (N-RGC) types, 5, 17, 30, and 40 (which comprised ISP 7^5^), matched the adult N-RGC types, with neonatal types 5 and 17 matching similarly the closely localized (**Fig. 4J**) adult types 26 and 29, and neonatal types 30 and 40 matching similarly the closely localized adult types 25 and 35 (**Fig. 4K**). The remaining adult N-RGC types 19 and 39 were also closely localized, and matched similarly neonatal Neurod2-expressing RGC types 10 and 28 (**Fig. 4L**).

**Figure 4.**
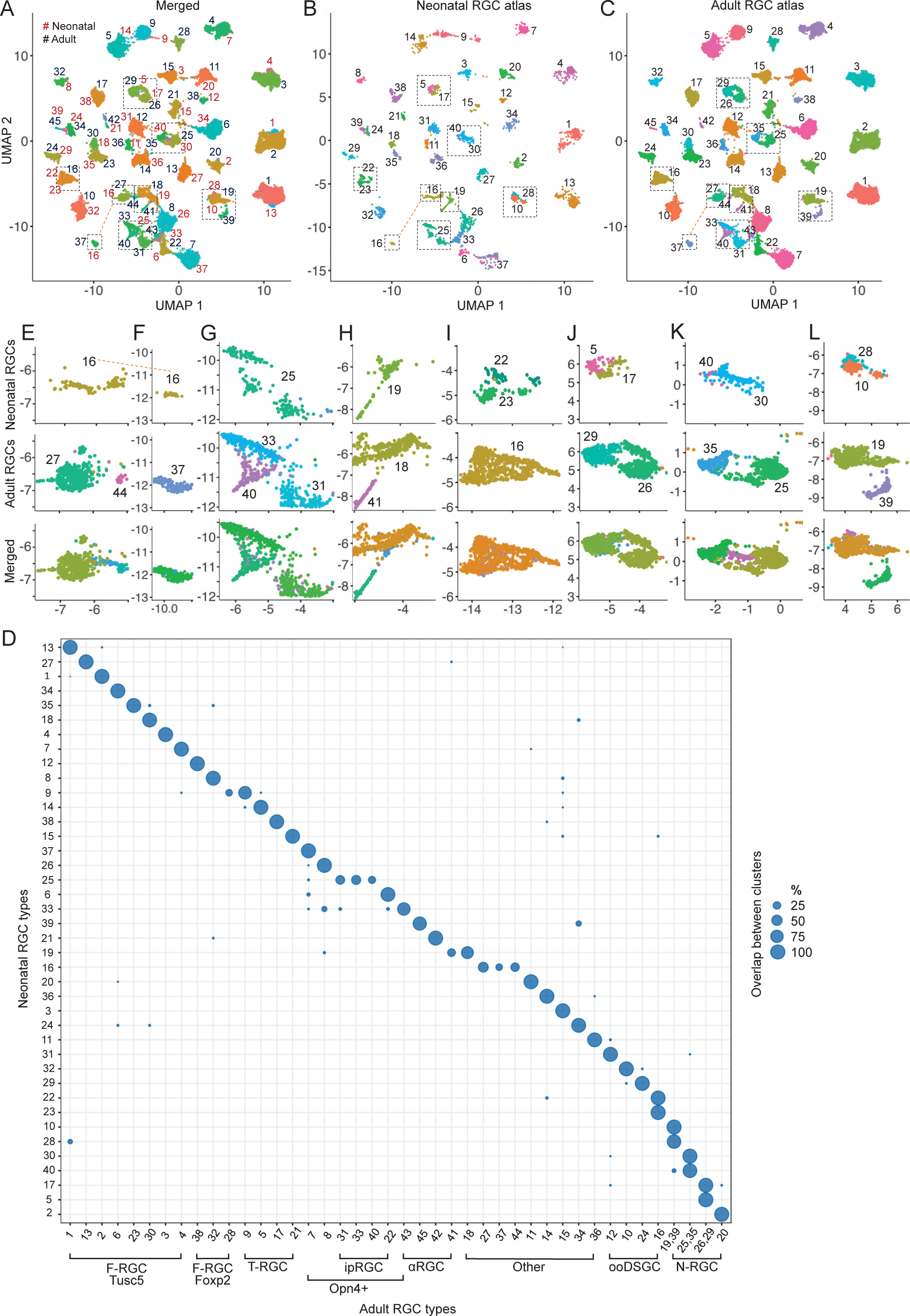
Neonatal RGC type origin of the mature adult RGC types. (**A-C**) Clustering together the neonatal and adult RGC datasets matched the different age types, as shown in the merged UMAP with neonatal RGC types numerated in brown font digits and adult RGC types numerated in black font digits (***A***). Neonatal RGC types from the merged UMAP (in *A*) are shown without the adult RGCs (***B***). Adult RGC types from the merged UMAP (in *A*) are shown without the neonatal RGCs (***C***). (**D**) Cluster-by-cluster percent overlap between the cells comprising neonatal and adult RGC clusters per the merged UMAP (in *A*). Bubble size scale bar indicates percent overlap. Broad RGC class annotation under the *x*-axis is per Table S2 in Tran et al., 2019^4^. (**E-L**) Insets (outlined in dashed-line rectangles in *A*-*C*) show instances of diverging or converging neonatal and adult RGC clusters per plot in panel *D*.

### Neonatal and adult RGC type markers

Because some of the gene-markers of RGC types change expression developmentally, we compared expression of the markers identified for neonatal and adult RGC types, and we also applied the algorithm we developed for determining unique single gene markers for individual RGC clusters^5^ to identify more single gene markers enriched in the adult RGC clusters. In our classification of neonatal RGC types, we identified several enriched single gene markers for each of the 40 clusters^5^. However, in the original analysis of the adult RGC clusters, 32 out of 45 clusters required combinations of enriched/unenriched genes to mark them uniquely, as single gene cluster markers were identified for only 13 clusters (see Fig. 1F and Supplementary Fig. 1F in Tran et al., 2019^4^). Of these 13 single gene cluster markers, 9 were amongst the neonatal RGC single gene markers we predicted previously (see Fig. 4 and Supplemental Data/Table 1 in Rheaume et al., 2018^5^; **Supplemental Table 1**). Of the remaining 4 genes, Ceacam10, Nmb, and Slc17a7, were developmentally upregulated from neonatal to adult RGCs (thus not sufficiently enriched in neonatal RGC clusters), and Rhox5 was enriched in several clusters of neonatal RGCs (and thus did not qualify as a neonatal cluster marker; **Supplemental Figure 6**) but was downregulated during maturation in these clusters, except for in cluster C32 (for which Rhox5 became a marker in adult RGCs).

Using the algorithm we developed for identifying single gene markers of neonatal RGC clusters^5^, we found single gene markers uniquely enriched in all adult atlas RGC clusters (**Supplemental Figure 7** and **Supplemental Data 1**). While all the single gene markers were statistically significantly enriched in the respective RGC clusters, and thus may point to the molecular pathways involved in the function unique to respective RGC subtype, not all of them are optimal candidates for labeling (e.g., by IHC, ISH, FISH) RGC types in retinal tissues, as some markers have a modest extent of enrichment or a subset of cluster cells did not expressed a respective marker (which may be related to a gene dropout per cell limitation in scRNA-seq methods^33^). Therefore, for labeling RGC types in retinal tissues, we selected only the most enriched singe gene markers, and in 5 cases two-gene markers (which were identified by an algorithm we previously developed for identifying a combination of the transcription factors uniquely co-enriched in individual neonatal RGC clusters^5^), without the need for negative and three-gene markers that were necessary for some clusters in the original analysis by Tran et al., 2019^4^ (**Fig. 5A**). Furthermore, as some of the RGC type markers change expression developmentally or after injury, we showed markers’ expression in both neonatal and adult RGC types (**Fig. 5C,D,F**), along with expression fold-changes during developmental maturation (**Fig. 5E,G**) and after injury (**Fig. 5B**).

**Figure 5.**
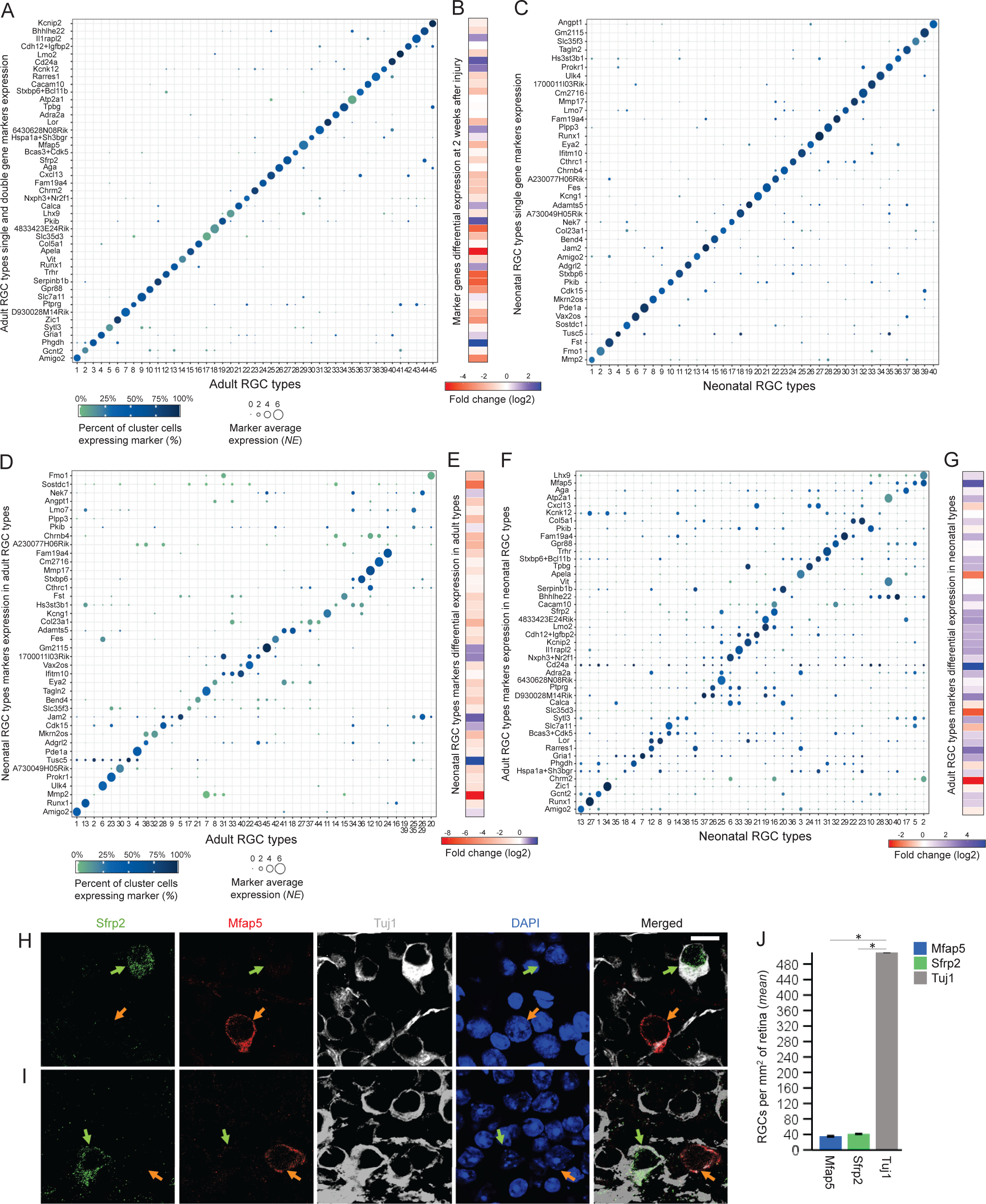
Neonatal and adult RGC atlas cluster markers enrichment and differential expression during maturation and after injury. (**A-B**) Bubble plot of positive single or double cluster marker genes enriched in adult atlas RGC types ≥1.5-fold relative to every other cluster. Bubble size (scalebar shown below the panel) indicates *z*-scores of average normalized gene expression in a cluster relative to all other clusters. For double markers, average of both markers’ *z*-scores was used. Average expression indicated by bubble size is for the mean of all the cells in a cluster. Bubble color (scalebar shown below the panel) indicates the percent of cells in a cluster for which expression of respective marker(s) was detected by drop-seq (***A***). Fold change of RGC type markers expression at 2 weeks after ONC relative to uninjured adult atlas RGC types (scalebar shown below the panel) (***B***). (**C**) Bubble plot of single cluster marker genes enriched in neonatal (P5) atlas RGC types ≥1.5-fold relative to every other cluster. Bubble size (scalebar same as for panel *A*) indicates *z*-scores of average normalized gene expression in a cluster relative to all other clusters. Average expression indicated by bubble size is for the mean of all the cells in a cluster. Bubble color (scalebar same as for panel *A*) indicates the percent of cells in a cluster for which expression of respective marker was detected by drop-seq. (**D-E**) Bubble plot of neonatal (P5) atlas RGC cluster markers expression in the adult atlas RGC clusters. Bubble size (scalebar shown below the panel) indicates *z*-scores of average normalized gene expression in a cluster relative to all other clusters. Average expression indicated by bubble size is for the mean of all the cells in a cluster. Bubble color (scalebar shown below the panel) indicates the percent of cells in a cluster for which expression of respective marker was detected by drop-seq (***D***). Fold change of neonatal atlas RGC type markers differential expression in adult atlas RGC types (scalebar shown below the panel) (***E***). (**F-G**) Bubble plot of adult atlas RGC cluster markers expression in neonatal (P5) atlas RGC clusters. Bubble size (scalebar same as for panel *D*) indicates *z*-scores of average normalized gene expression in a cluster relative to all other clusters. For double markers, average of both markers’ *z*-scores was used. Average expression indicated by bubble size is for the mean of all the cells in a cluster. Bubble color (scalebar same as for panel *D*) indicates the percent of cells in a cluster for which expression of respective marker(s) was detected by drop-seq (***F***). Fold change of adult atlas RGC type markers differential expression in neonatal atlas RGC types (scalebar shown below the panel) (***G***). (**H-I**) Immunohistological analysis of Sfrp2 and Mfap5 expression pattern in the retina is consistent with bioinformatic prediction of these genes labeling non-overlapping subpopulations of RGC types, C27 and C29, respectively (as shown in *A*). Confocal representative images of different regions (shown in ***H*** and ***I***) of flat-mounted adult retina immunostained for the predicted RGC type C27 marker Sfrp2, predicted RGC type C29 marker Mfap5, neuronal marker Tuj1, and nuclear marker DAPI. Green arrows point to the Sfrp2+/Tuj1+ RGCs, and orange arrows point to the Mfap5+/Tuj1+ RGCs. Mfap5-labled and Sfrp2-labled subpopulations of RGCs do not overlap, whereas most RGCs (Tuj1-labled cells) were not labeled by either marker. Scale bar: 10 μm. (**J**) Quantifications of Tuj1+/Mfap5+/Sfrp2-, Tuj1+/Mfap5-/Sfrp2+, and Tuj1+/Mfap5-/Sfrp2-cells in flat-mounted adult retinas; no Tuj1+/Mfap5+/Sfrp2+ cells were found (mean ± SEM shown, *n* = 4 cases per group). Data analyzed using ANOVA, overall *F* = 1469, *p* < 0.001, with *p*-values of pairwise comparisons determined by posthoc LSD. Significant differences (*p* < 0.001) indicated by an asterisk (*).

Several of the predicted marker genes have been previously validated. For example, we verified expression in the RGCs by immunostaining or FISH^5^ for: Neonatal cluster 34 marker Zic1, which is also most highly enriched in the matching adult cluster C6 compared to the other clusters (**Fig. 5A**); neonatal cluster 27 marker Runx1, which is also most highly enriched in the matching adult cluster C13 (**Fig. 5A**); and neonatal cluster 3 marker Fst, which is also enriched in the matching adult cluster C15 (**Supplemental Figure 7**). Also, previously validated^32^ neonatal αRGC cluster 39 marker Kcnip2 (see Supplemental Data in Rheaume et al., 2018^5^) was most highly enriched in the matching adult αRGC cluster C45 (**Fig. 5A**). More markers we previously predicted for the neonatal RGC clusters^5^, which were also enriched in the adult RGC clusters, were validated in adult RGCs (see Tran et al., 2019^4^). Here, we also validated by immunostaining in retinal tissues two of the new single gene markers we predicted for adult RGC clusters: Sfrp2 marker of uncharacterized Other type C27, and Mfap5 marker of N-RGC type C29 (**Fig. 5A**). We quantified Sfrp2+/Tuj1+ and Mfap5+/Tuj1+ RGCs as percent of total RGCs (Tuj1+) in retinal tissues, and found that they represent 1.73% and 1.48% of the RGCs, respectively, which is consistent with the Sfrp2-labled C27 representing 1.41% and Mfap5-labled C29 representing 1.38% of all RGCs, respectively, based on the scRNA-seq cluster markers analysis (**Fig. 5H-J**).

### Global properties of the transcriptome predict the resilience to injury of two **_α_**RGC types

To gain insight into the global transcriptomic dynamics of RGC types, we analyzed whether the extent of similarly/dissimilarity between RGC types changes during maturation and after injury. We found that the dissimilarity between the transcriptomes of adult atlas RGC types is within an average *r* range of 0.91-0.97 (Pearson; **Fig. 6A**; even if the atlas RGCs are downsampled to match the number of injured RGCs in the analysis above; **Supplemental Figure 8**), which is twice the range we found for pre-eye-opening neonatal atlas RGC types (0.96-0.99 average *r* range^5^). Furthermore, the extent of dissimilarity between the transcriptomes of RGC types also increased after injury to 0.59-0.95 average *r* range (Pearson; **Fig. 6B**). However, there was no significant correlation between survivability and the mean transcriptome size of uninjured (*r* = 0.26, *p* = 0.07) or injured (*r* = 0.05, *p* = 0.74) RGC types (**Supplemental Figure 5B-C**), nor the relative proportion of mitochondria-related genes (see above) of injured types (*r* = 0.17, *p* = 0.25; **Supplemental Figure 5A**). We also showed that the varying proportion of highly expressed (>0.5 *NE*) genes^5,34^ was modestly correlated between uninjured and injured RGC types (*r* = 0.31, *p* < 0.05; **Fig. 6C-F**), but it was not correlated to RGC survivability (*r* = −0.05, *p* = 0.73), suggesting that this varying global transcriptomic property of RGC types is overall preserved after injury in most RGC types, regardless of susceptibility or resilience of an RGC type. Finally, we found that the differences between the transcriptomes of uninjured RGC types (average cluster *r* values in **Fig. 6A**) is significantly correlated to the survivability of injured RGC types (*r* = −0.44, *p* < 0.005; **Fig. 6G**), and that the types with the most influential regression residuals accounting for correlation are the resilient atlas types (C42 and C45) belonging to the αRGC class (see Table S2 in Tran et al., 2019^4^; **Fig. 6G-H**).

**Figure 6.**
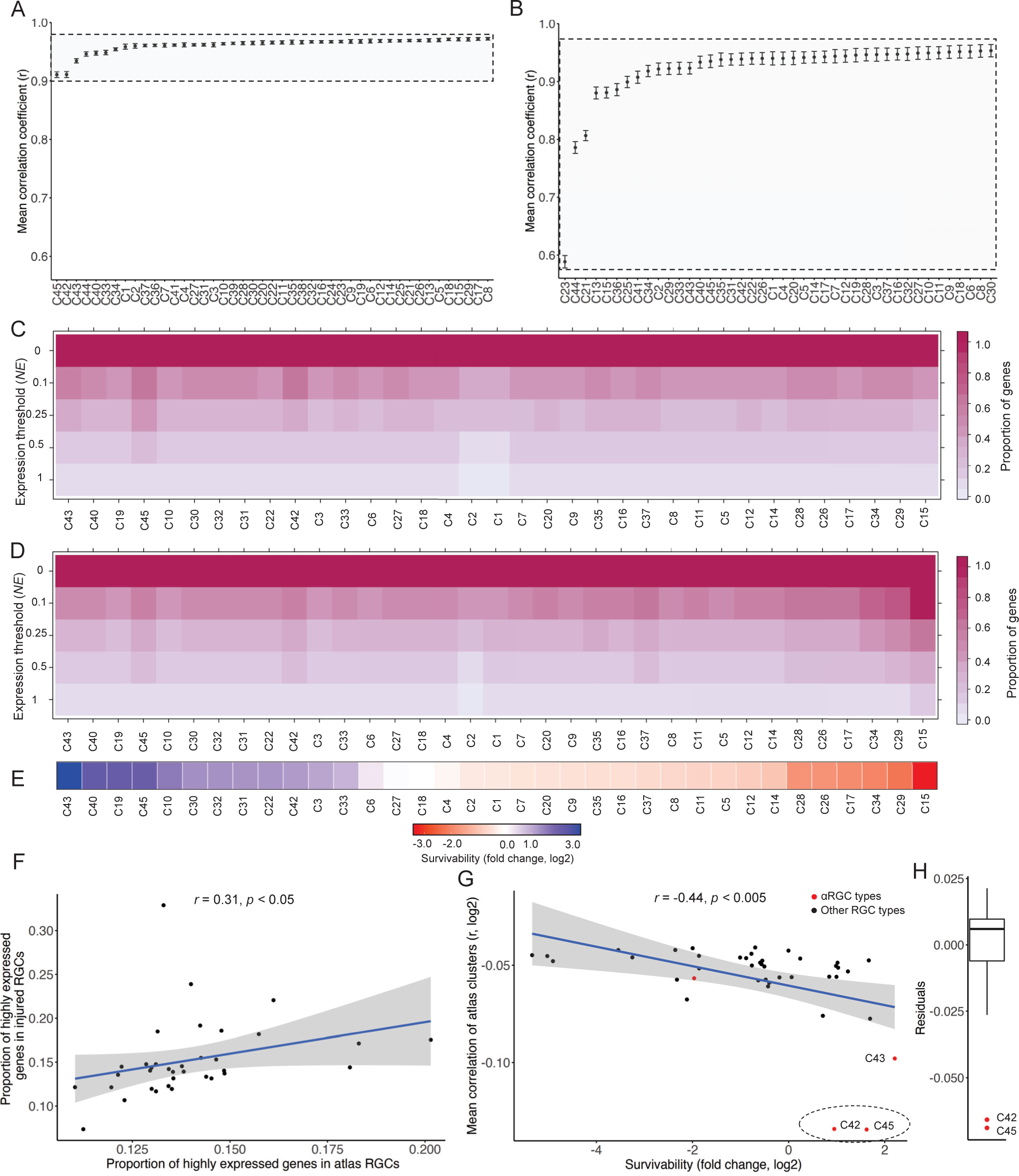
The effects of injury on global transcriptome parameters of RGC types. (**A-B**) Mean correlation coefficient (*r*) of atlas RGC clusters compared to every other uninjured atlas RGC cluster (***A***) and of injured RGC clusters compared to every other injured RGC cluster (***B***; at 2 weeks after ONC). The grey area (in *A* and *B*) represent the inter-cluster variance of Pearson correlation coefficients, highlighting >2-fold increase in variability of RGC transcriptomes after injury. C24, C38, and C39, which did not survive 2 weeks after injury, are not shown in *B*. Error bars = SEM. (**C-D**) Proportions of low and highly expressed genes at different *NE* ranges, as marked, in uninjured atlas (***C***) and injured (***D***) RGC clusters. The order of the clusters (left to right) is the same in *C*-*E*. Clusters that contain 0 or < 5 cells at 2 weeks after ONC are excluded from analyses in *C*-*F*. (**E**) Survivability of RGC clusters (at 2 weeks after ONC), based on fold change (log2) of uninjured atlas to injured RGC clusters. (**F**) Linear regression of the proportion of highly expressed genes (>0.5 *NE*) in uninjured atlas RGC clusters vs. the proportion of highly expressed genes (>0.5 *NE*) in injured (2 weeks after ONC) RGC clusters. Gray area around the fit-line (in *F* and *G*) indicates 95% confidence interval for the linear regression model. (**G**) Linear regression of mean correlation coefficients of uninjured atlas RGC clusters compared to every other cluster (shown in *A*) vs. survivability of injured RGC clusters (ranked in *E*; at 2 weeks after ONC). The red clusters belong to the αRGC class, as marked. (**H**) A boxplot of the residuals for the regression in (*G*), with the most influential datapoints (i.e., outlier residuals shown in red) corresponding to the αRGC clusters that are shown in dashed shaded oval in *G*.

## DISCUSSION

The CellTools algorithms we developed for denoising the reference dataset and machine learning-optimization of the *k*-weight parameter, improved the integration mapping of query-to-reference scRNA-seq datasets, thereby addressing the need in the field for new advancements of integration mapping algorithms^1-3^. Retrospect analyses of the noise cells detected by our DeNoise algorithm showed that they had either abnormally large transcriptomes (typical when a piece of a broken, or another cell, is attached to a bead with an intact cell) or an abnormally high proportion of mitochondrial genes (typical for cells progressing towards apoptosis), were not overrepresented in any clusters, and their removal improved the accuracy of integration. Moreover, our approach enabled removing sources of noise not detected by a doublet-removal algorithm (Scrublet^24^). Further improvement in the accuracy of cell-to-cluster assignment in the integration mapping was achieved by the machine learning algorithm we developed for choosing the *k*-weight parameter of the KNN algorithm in a data-driven manner. Our two-step algorithm outperformed other integration mapping algorithms on comparative analysis, and reduced incorrect integration mapping of the query cells with the reference dataset by nearly 4-fold, thereby resetting the bar in the field from several percent of incorrect integration to just under one percent. Because we also found that the denoising step of our algorithm improved the performance of other integration mapping algorithms (Seurat’s MapQuery^17^ and Symphony^27^), it is possible that through further development the DeNoise algorithm may be adopted for application in the non-mapping integration tools as well, such as those that were designed for batch adjustment (e.g., RPCI^35^, mtSC^36^) or for matching scRNA-seq datasets with other types of corresponding single cell datasets (e.g., by GLUE^37^ for matching with scATAC-seq, snmC-seq3, sci-MET4 datasets).

As changes in gene expression during various biological and pathological processes require integration mapping algorithms for tracing respective cell type origin, our algorithm will assist studies across biological fields aimed at tracing cell type-origin of injured, diseased, developing, matured, or otherwise broadly-changed transcriptomes. As a proof-of-concept, we showed that our algorithm improved the identification of scRNA-seq-derived resilient and susceptible RGC types, that were originally classified using the algorithm iGraphBoost^4^. We found that the originally unassigned injured RGCs were not substantially different from other injured RGCs in any of the global transcriptomic properties, and that RGC types C2, C19, C20, C26, C32, C35, C37, C42, and C45 were substantially overrepresented amongst the originally unassigned injured RGCs. Consequently, the re-analysis of the RGC types’ survivability amended identification of the originally reported resilient and susceptible RGC types^4^. For example, αRGC types were previously shown to be more resilient to injury^28^ and, along with the ipRGC types, more responsive to Pten inhibition for regenerating axons^12,13,29^. An αRGC type C43 was indeed the most resilient type in the Tran at el. (2019) analysis, and it remained in that rank after we assigned the originally unassigned RGCs to types. However, an αRGC type C45 was ranked as susceptible in the Tran at el. (2019) analysis, but it changed to resilient after we assigned the originally unassigned RGCs to types. This change in classification of an αRGC type C45 from susceptible to resilient by our analysis is consistent with previous studies that showed that αRGC types are more resilient to injury than other RGC types. This finding further supports the improved accuracy of our algorithm. Thus, the updated ranking of RGC type resilience presented herein, will assist the studies that rely on RGC type identification as resilient or susceptible in order to investigate the differences in their underlying biology and to identify neuroprotective and risk genes for therapeutic targeting.

We then used algorithms we developed previously^5^, to identify single and double gene markers uniquely enriched in the adult atlas RGC clusters^4^, which may be used for labeling RGC types by IHC, ISH, FISH in retinal tissues. We found single gene markers for most clusters and two-gene (all positive) markers for other clusters, in contrast to the majority of clusters requiring combinations of two or three positive and negative markers in the original analysis by Tran et al., 2019^4^. We also provided a heatmap of statistically significantly co-enriched cluster-specific single genes, which may point to the molecular pathways involved in the function unique to respective RGC type, although they are not optimal candidates for labeling RGC types in retinal tissues (in contrast to the single and double gene markers described above), as some of these genes have only modest extent of enrichment or a portion of cluster cells did not express a respective marker (which may be related to a gene dropout per cell limitation in scRNA-seq methods^33^).

We also found that many of the adult RGC type markers were uniquely enriched in the developing neonatal atlas RGC clusters too^5^, and some have been previously validated (e.g., Pde1a, Zic1, Kcnip2) as RGC type-specific^4,5^. We then validated by immunostaining two of the new single gene markers we predicted for adult RGC clusters: Sfrp2 marker of uncharacterized Other type C27 and Mfap5 marker of N-RGC type C29, which labeled in the retinal tissues similar proportions of the RGCs as was predicted based on the scRNA-seq analysis. Because we also found that some markers were cluster-specific only in adult RGCs, as during maturation from the neonatal RGCs they were either upregulated (e.g., Ceacam10 for C37, Nmb for C40, Slc17a7 for C25) or downregulated (e.g., Rhox5 for C32), we characterized the correspondence between the adult and neonatal RGC types (after applying DeNoise to the neonatal RGC reference dataset too, which identified ∼2.9% of cells as noise). We found that while most types matched one-to-one, others diverged or converged into a few clusters during maturation. We also analyzed expression profile of the neonatal RGC types markers in adult RGC types, and vice versa, as well as adult RGC type markers expression profile at two weeks after ONC, and showed fold-changes for markers which were upregulated or downregulated.

Additionally, we provided insights into the global characteristics of RGC types^5,34^ and how axonal injury affects them. We showed that dissimilarity between RGC types’ transcriptomes, although modest, increases over two-fold during developmental maturation, which may be related to the fact that there are more types of RGCs in adult (46) than there are in neonatal (40)^5^, as the cells continued to diverge further towards their final state during maturation into adulthood^4^. We then found that the most divergent atlas RGC types are resilient and belong to the αRGC class, while after injury dissimilarity between the transcriptomes of RGC types increased overall. However, the proportion of highly expressed genes^34^, another varying global property of the RGC types transcriptome^5^, was overall preserved after injury in most RGC types regardless of susceptibility or resilience of an RGC type. We also showed that neither the cluster size (average number of cells), cluster transcriptome size (average number of expressed genes), nor the proportion of cellular stress genes^1,22,23^ were significantly correlated to the differential survivability between RGC types. Thus, while the factors unique to susceptible or resilient RGC types are primary determinants of survivability and may be associated with increases in dissimilarity between the transcriptomes of RGC types after injury, the differences between global properties of atlas RGC transcriptomes are associated with the survivability of at least two αRGC types. This is the first time a global parameter of an uninjured cell type transcriptome has been linked to a neuronal response to injury, suggesting that transcriptomic global parameters are important to consider in future studies.

We also built the Subtypes Gene Browser for cluster-by-cluster differential gene expression analysis between the atlas and injured RGC clusters (https://health.uconn.edu/neuroregeneration-lab/subtypes-gene-browser). Finally, we made the algorithms we developed publicly available through the R-package, CellTools, which will assist scRNA-seq studies across biological fields (see Availability and Implementation in the Methods section).

## SUPPLEMENTARY INFORMATION

Supplementary Information includes 8 Figures, 1 Table, and 5 Data files.

## ACKNOWLEDGMENTS

This work was supported by grants from The University of Connecticut School of Medicine, Start-Up Funds (to E.F.T.), the BrightFocus Foundation (Grant G2017204, to E.F.T.), and the National Institutes of Health (NIH) (Grant R01-EY029739, to E.F.T.). Portions of this research were conducted at the High Performance Computing Facility, University of Connecticut. We thank Nicholas Tran, Karthik Shekhar, and Joshua Sanes (Center for Brain Science and Department of Molecular and Cellular Biology, Harvard University, Cambridge, MA, Broad Institute of Harvard and MIT, Cambridge, MA, University of California, Berkeley CA, and Baylor College of Medicine, Houston TX) for providing us with raw data from which they generated the figures and tables in Tran et al. (2019). We also thank Sophan Iv and Vijender Singh (Research IT Services, University of Connecticut), and Stephen King (High Performance Computing Facility, University of Connecticut), for assistance with computational resources and the development of the website. Finally, we thank Ashiti Damania and Mahit Gupta (undergraduate students, University of Connecticut) for technical assistance.

## AUTHOR CONTRIBUTIONS

B.A.R. performed the analysis and contributed to conceptualization of the study, J.X. performed the analysis, W.C.T and S.D. assisted with the analysis, and E.F.T. conceptualized the study and wrote the manuscript.

## METHODS

### Animals

All animal procedures were approved by the University of Connecticut Institutional Animal Care and Use Committee and by the Institutional Biosafety Committee at the University of Connecticut, and performed in accordance with the ARVO Statement for the Use of Animals in Ophthalmic and Visual Research. C57BL/6J mice were obtained from Charles River Laboratories, Inc.

### Histological procedures and quantifications

Standard histological procedures were used, as described previously^5,16,38^. Briefly, 10-week-old mice were anesthetized and transcardially perfused with 0.9% saline solution followed by 4% paraformaldehyde (PFA). The eyes were resected and post-fixed 2 hours at room temperature. The retinas were then dissected-out from the eyes in PBS, and after making 4 symmetrical slits flattened retinas were embedded in OCT Tissue Tek Medium (Sakura Finetek), frozen, and cryostat-sectioned at 14 µm horizontality (capturing the ganglion cell layer of the whole retina) onto Superfrost Plus glass slides (VWR International, LLC). For immunostaining, the cryosections were blocked with appropriate sera, incubated overnight at 4 °C with primary anti-Sfrp2 (1:250; mouse monoclonal, Sc-365524, SCBT), anti-Mfap5 (1:300; rabbit polyclonal, ab232846, Abcam), and anti-βIII-Tubulin (Tuj1; 1:500; rabbit polyclonal, ab18207, Abcam) antibodies, counterstained with DAPI (1:5000; Thermo Fisher Scientific), then washed three times, incubated with appropriate fluorescent dye-conjugated secondary antibodies (1:500; Alexa Fluor, Thermo Fisher Scientific) overnight at 4 °C, washed three times again, and mounted for imaging. Images were acquired using confocal microscope (40x/1.3 Achrostigmat Oil; Zeiss Confocal, LSM 880). For quantification, Mfap5+/Tuj1+ and Sfrp2+/Tuj1+ cells were counted, using ImageJ software, as percent of total Tuj1+ cells per mm^2^ in dorsal, ventral, temporal, and nasal regions of the retina, and then averaged. Total of four retinas were quantified to estimate overall percent of the Mfap5-labled and Sfrp2-labled RGCs per mm^2^ of the retina. No doble positive, Mfap5+/Sfrp2+, cells were found.

### Single cell RNA-seq data procurement and initialization

BAM files, raw counts, normalized matrices, and cell metadata (e.g., type assignment) for the mouse adult RGC atlas and the injured RGCs were obtained from the Gene Expression Omnibus (GEO) accession number GSE137400^4^. Atlas and injured RGC BAM files were converted to FASTQ files using CellRanger’s bamtofastq software. FASTQ files from both the atlas and injured datasets were aggregated where appropriate using CellRanger and then mapped to the CellRanger mm10-1.2.0 transcriptome, as the original analysis had mapped the atlas and injured data to different transcriptomes, making comparisons between them more difficult and, in some cases, impossible.

Because the RGCs were sequenced in separate batches, we first performed batch correction using the Seurat’s v. 4.0.3^1,17^ FindIntegrationAnchors and IntegrateData functions. Normalization of the raw counts was performed using the Seurat function, NormalizeData, which divides the feature counts by the number of counts per each cell and then applies natural log transformation^1,17^. The same cells that passed the initial quality check and thresholding, and the same type assignments for the atlas, from the original analysis^4^ were used in the downstream characterization and analysis by the present study.

### Dimensionality reduction

Dimensionality reduction, visualization, and model generation for the subsequent reference mapping was performed using the uniform manifold approximation and projection (UMAP) implementation^18,19^ in Seurat v. 4.0.3 using default parameters^1,17^. Feature selection for the UMAP algorithm was determined using the top 50 dimensions (principal components; PCs) for atlas, and 30 dimensions for injured, RGCs, which were respectively determined to be significant sources of variation by Seurat’s JackStraw implementation^17,39^. UMAP of atlas RGCs were color-coded with their original cluster designations^4^ for determining the cluster-latent-space relationship.

### Denoising and reference-mapping of query cells using CellTools’ DeNoise and MapTo

First, the DeNoise algorithm removes the sources of noise from the reference UMAP by identifying outlier cells in the cluster-UMAP relationship. To do this, each cluster’s quartiles and interquartile range (IQR) is calculated for the x and y dimensions of the UMAP. Any cell *i* belonging to cluster *c* (*c*,*i*) whose x or y dimensions are more than 1.5 times the IQR, either greater than the upper quartile (Q3) or less than the lower quartile (Q1), is considered to be *noise* within the UMAP structure and is subsequently removed for downstream analyses. The following equations summarize the discovery of *noise* and *signal* cells:

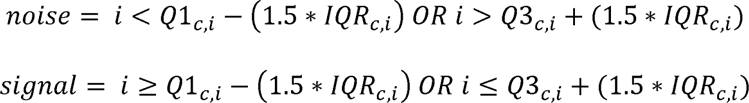

After *noise* cells are removed from the original UMAP, a new UMAP is generated with only the *signal* cells using Seurat’s RunPCA and RunUMAP (with return.model = T) functions. Query cells transcriptomes are then mapped to the denoised reference UMAP model using the MapTo function from CellTools, which we developed (using the top 50 PCs for RGCs, and see the parameters used for other cell types in Supplemental Data files 2-5). This second step of our algorithm is a machine learning function, MapTo, which utilizes a wrapper/iterative-looper for Seurat’s FindTransferAnchors, TransferData, IntegrateEmbeddings, and ProjectUMAP functions^17^. It first identifies the optimal *k*-weight parameter for the KNN latent search-space by iteratively mapping a test-query dataset to a test-reference dataset with known cell types. To accomplish this, the algorithm first randomly subsamples the reference dataset into a test-query dataset. By default, 20% of the cells are subsampled, but this parameter can be changed by user (for the RGCs we used 23.7%, or 8,456 cells, which matched the size of the query dataset). The remaining cells from the reference dataset are designated as the test-reference dataset, and the test-query dataset is iteratively mapped to the test-reference dataset using every *k*-weight parameter within the user-defined range (default is between 10-100; see **Supplemental Figure 1B**). Iterations are then ranked based on the percent of correctly mapped cells, and the iteration with the highest percent of accurately mapped cells is selected as the optimal *k*-weight parameter (if more than one *k*-parameter yields same percent of accurately mapped cells, then the smaller *k*-parameter is selected; for this study the optimal *k* = 23). Next, the query dataset is mapped to the denoised reference dataset using the optimal *k*-weight parameter. The predicted cell IDs (i.e., the reference dataset IDs transferred to the query data; RGC types here) are outputted in the returned query Seurat object, along with a reference-based UMAP of the query cells.

### Single cell RNA-seq datasets of different cell types used for comparative analysis of integration mapping algorithms

The raw scRNA-seq dataset of adult atlas mouse RGCs (*n* = 34,584), which was used herein in the initial validation of the 2-step (MapTo and DeNoise) integration mapping algorithm, is available at the NCBI GEO under accession number GSE137400^4^. The raw scRNA-seq dataset of adult human peripheral blood mononuclear cells (PBMC) (*n* = 13,999), which was used in validation of the Seurat’s MapQuery integration mapping algorithm^17^, is available at the NCBI GEO under accession number GSE96583^40^. The raw scRNA-seq dataset of adult human SARS-CoV-2-infected PBMCs (*n* = 8,850), which was used in validation of the Symphony integration mapping algorithm^27^, is available at the NCBI GEO under accession number GSE149689^41^. The raw scRNA-seq dataset of adult mouse and human pancreatic islets cells (PIC) (*n* = 12,474), which was used in validation of the meta-clustering method asCIDER (CIDER) integration mapping algorithm^25^, is available at the NCBI GEO under accession number GSE84133^42^. The raw scRNA-seq dataset of embryonic mouse pancreatic epithelial cells (PEC) from pancreatic endocrinogenesis (*n*L=L22,163), which was used in validation of the semi-supervised integration scANVI (scArches) integration mapping algorithm^26^, is available at the NCBI GEO under accession number GSE132188^43^.

### Integration merging of the neonatal RGC atlas and adult RGC atlas

Seurat (described above^17^) objects for clusters of the mouse neonatal RGCs (NCBI GEO under accession number GSE115404^5^) and adult RGCs (described above, NCBI GEO under accession number GSE137400^4^) were merged. Then, Monocle v.3^44^ was used to group cells into clusters based on their gene expression profiles.

**Availability and implementation** (design of the website and online tools). All R codes and scripts designed for this study will be shared upon request. The R and python scripts used for implementing mapping integration software in the comparative analysis are in the Supplementary Data files 2-5. The Subtypes Gene Browser was designed in the same format as we previously did for RGC subtypes^5^, using R and ShinyApps with R-markdown language^45^. Boxplots, violin plots, and bar plots were adapted from ggplot2 R software package for data visualization^46^. CellTools is a suite of single-cell analytical tools that was developed for this study and others, and is available at https://health.uconn.edu/neuroregeneration-lab/CellTools.

### Data availability

The raw and processed data used in this study are available through the NCBI GEO under accession numbers GSE137400^4^ and GSE115404^5^. Data processed in this study are available through the NCBI GEO under accession number GSE205135. Access to the dataset is also available through a user-friendly Subtypes Gene Browser web application, https://health.uconn.edu/neuroregeneration-lab/subtypes-gene-browser.

### Thresholds for signature genes enriched in adult atlas RGC types

For determining unique single gene markers enriched in individual adult atlas RGC clusters (shown in **Supplemental Figure 7**), we applied an algorithm we described previously in the classification of neonatal RGC types^5^, using the following thresholds. Genes uniquely enriched per cluster were expressed (>0.05 *NE*) at least 1.8-fold more compared to every other cluster, respectively, at *p*-valueL≤L0.05. The *p*-value had to pass ≤L0.05 on two separate tests: independent samples *t*-test (2-tailed) and a nonparametric independent samples Mann–Whitney *U* test (using R software). Enriched gene markers that didn’t pass one or both statistical tests but met the first two criteria are listed in **Supplemental Data 1**. The threshold of 1.8-fold was selected because it predicted at least one uniquely enriched gene marker for all clusters. To identify a combination of genes to mark clusters in **Fig. 5A**, we applied an algorithm we developed previously for identifying a combination of transcriptional regulators uniquely co-enriched in a cluster^5^.

### Bubble plots, Violin plots, Density plots, Heatmaps, and Boxplot

The bubble plots (for **Figs. 4-5** and **Supplemental Figure 4**) were generated using the Ggplot2 geom_point() R function. The heatmap for **Supplemental Figure 7** was generated using the R package Superheat^47^; expression values for each gene across different clusters were normalized with *z*-scores for each row using the centered Scale R function prior to plotting, and the columns were then specified in an increasing order from cluster 1 through 45. Violin plots were generated using Seurat’s VlnPlot function. Base R statistical software functions were used for generating the density plots (density function from the stats package) and boxplot (boxplot function from the graphics package).

### CellTools’ GlobalProbe

We designed the function GlobalProbe to explore the relationship between the global transcriptomic properties and cell types, and to test whether these properties can predict (using a linear model) cell type-specific phenotypes (e.g., resilience to injury, which is a dependent variable here). The function requires a Seurat object as input and the designation of the cluster label slot by which the user intends to make comparisons. The function will output the mean and SEM of correlation coefficients (Pearson’s) for the comparisons between each cluster’s mean gene expression and every other cluster’s mean gene expression. The outputted graph is formatted as in **Fig. 6A-B**. If a dependent variable for a cell type-specific phenotype is passed to the function (by default this option is off), the function will output the graph formatted as in **Fig. 6G**.

The model for these data is as follows: *f*(*x*) = −0.0038x - 0.06 This linear model and its residuals (outliers shown **Fig. 6H**) demonstrate that global transcriptomic properties can predict cell type-specific phenotypes, which would be indicated by a significant (*p* < 0.05) coefficient of determination (*R*^2^).

### Statistical analyses

Pearson (2-tailed) correlation analyses were used for the transcriptomes of cells and clusters, and the mean correlation coefficient *r* was computed as reported, with mean ± SEM shown and significance cutoff at *p* ≤ 0.05. In **Fig. 6F-G** and **Supplemental Figure 3B**, correlations were visualized using linear regressions. For **Fig. 6F**, clusters which had no cells that survived 2 weeks after ONC were excluded from the correlation and regression analyses. For **Supplemental Figure 3A**, analysis of significance was performed using ANOVA with posthoc LSD (SPSS). Fold change significance in Fig. 3B was determined using the EdgeR algorithm^48^.

