## Supplementary Information for "CellTools algorithm for mapping scRNA-seq query cells to the reference dataset improves the classification of resilient and susceptible retinal ganglion cell types"

Rheaume, et al.

### Supplementary Information

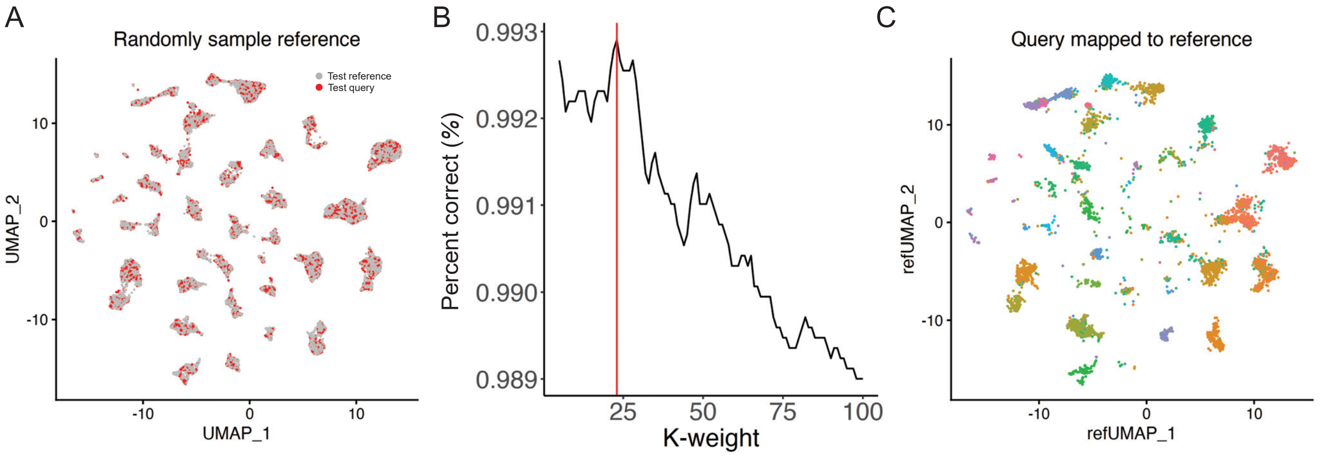


**Supplemental Figure 1. Steps MapTo algorithm performs to optimize the *k*-weight parameter**

(**A**) After the reference dataset (adult RGC atlas here) is denoised by the DeNoise algorithm, 20% of the cells are randomly subsampled and set aside as a “test query” dataset (red cells), and the remainder of the cells become the “test reference” dataset (gray cells). The cells are represented on the denoised reference dataset (atlas UMAP in Fig. 1C), with the “test query” (red cells) shown in the original reference clusters from which they were subsampled.

(**B-C**) The “test query” dataset cells iteratively mapped over a range of *k*-weights to the “test reference” dataset, and the *k*-weight that yields the highest percent of correctly mapped cells is selected as optimal for mapping the original query dataset. Here, $k$ = 23 (red line) results in 99.3% of the cells being correctly mapped to their cluster-origin (from which they were originally subsampled) in the reference dataset (***B***), as visualized on the denoised reference dataset (atlas UMAP in Fig. 1C) structure (color-coded as the original clusters, see clusters’ colors annotation in Fig. 1D), with only the mapped “test query” cells shown (***C***).

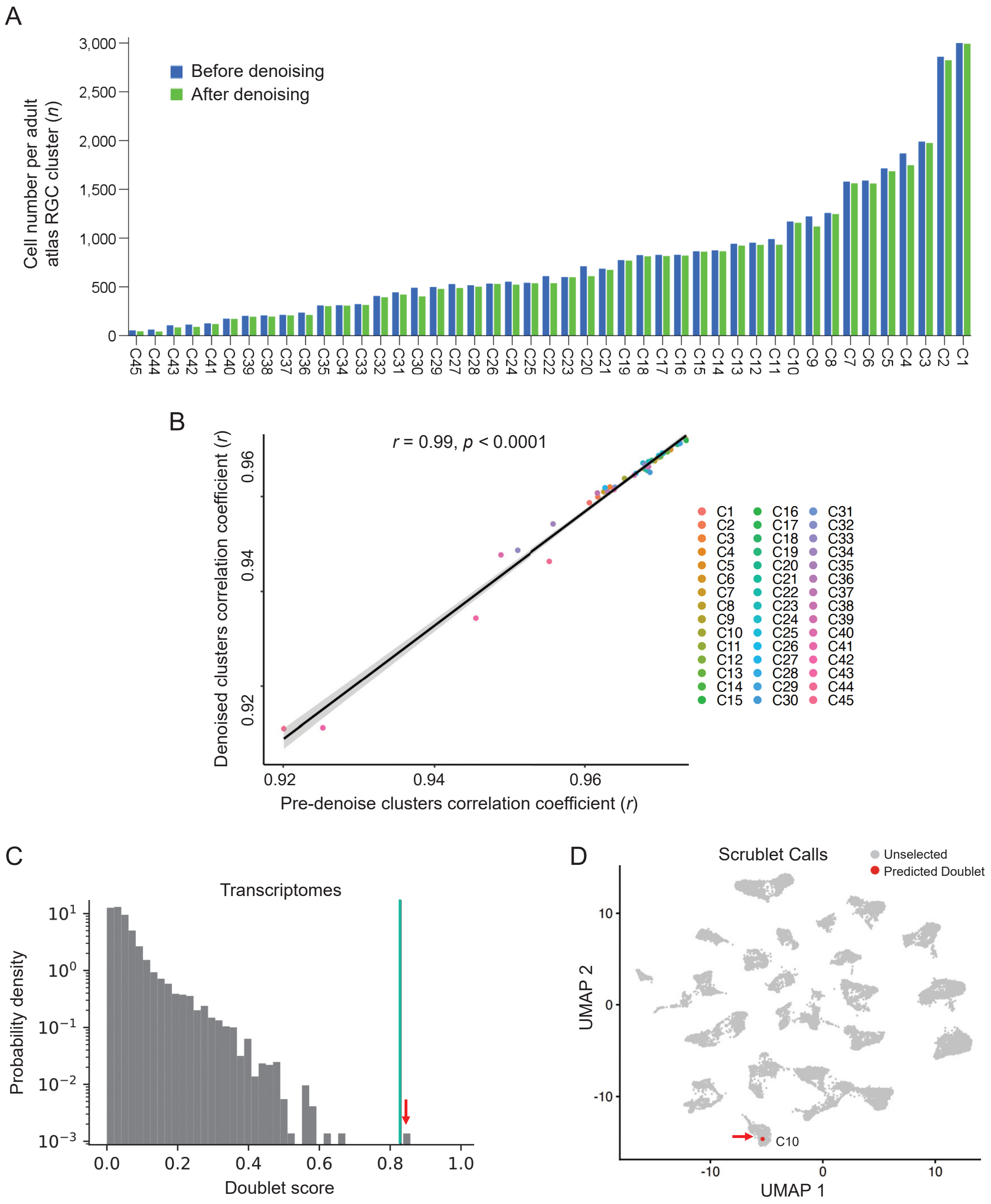


**Supplemental Figure 2. Noise cells proportions and potential doublets in atlas RGC clusters**

(**A**) Comparisons of cell numbers for each adult atlas RGC cluster before and after removal of the noise cells.

(**B**) Average correlation between the clusters before and after denoising is high ($r$ = 0.99). An $r$-value is also high ($r$ = 0.99) for the shown here linear regression of mean Pearson's correlation coefficients of the pre-denoise clusters compared to every other pre-denoise atlas RGC cluster vs. mean Pearson's correlation coefficients of the denoise clusters compared to every other denoise atlas RGC cluster.

(**C**) Predicted doublet score probability distribution plotted as a histogram of the transcriptomes (plot generated by Scrublet). Green line indicates Scrublet-predicted score threshold for a positive doublet. Red arrow indicates the only one transcriptome/cell that slightly exceeded this threshold and is a predicted doublet.

(**D**) The UMAP reference of uninjured adult atlas RGCs (from Fig. 1B) prior to denoising, with a red arrow indicating the Scrublet-predicted doublet, which did not present as an outlier in the post-clustering structure of the dataset and therefore was not amongst the "noise" cells detected by our algorithm. Mean UMI count per cell in the cluster C10 (to which this cell belongs) is 10,224 and this transcriptome/cell had 13,026 UMI counts that fell within 1 SD of the C10 mean (SD = 0.58), therefore, it is unlikely to be a doublet.

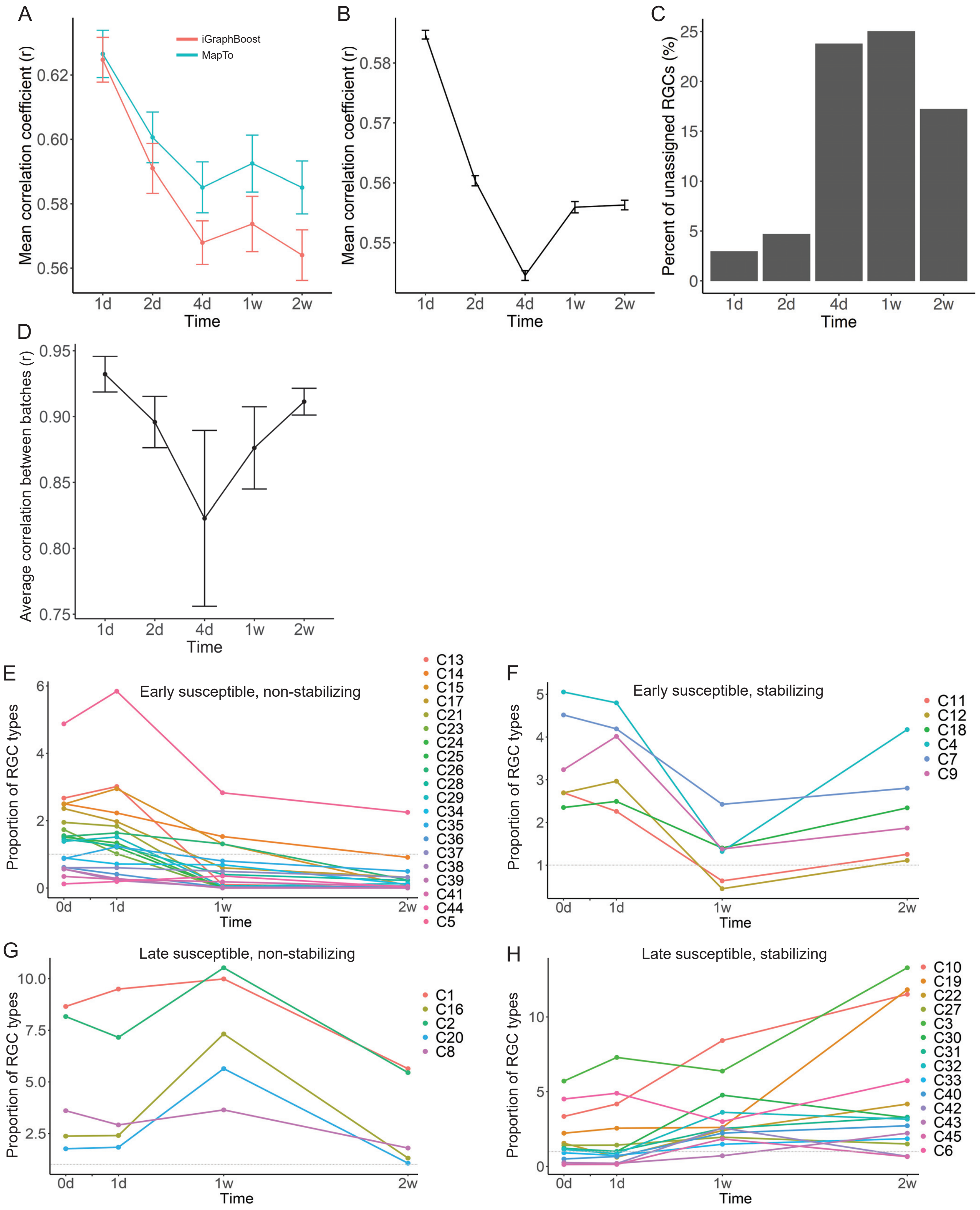


**Supplemental Figure 3. Changes in RGC type transcriptomes over time after injury are non-linear**

(**A**) Mean correlation coefficients (*r*), at different timepoints after injury, between the transcriptomes of uninjured atlas RGC clusters and the corresponding injured RGC clusters assigned by the iGraphBoost (red line) or two-step MapTo (green line) algorithms, show significantly higher mean *r* for MapTo assignments. Significant main effect difference between the algorithms was determined by ANOVA ($F = 7.6$, $p < 0.01$; there was no interaction effect with the time variable). Error bars = SEM.

*See continuation of the Supplemental Figure 3 legend on the next page.*

***Continuation of the Supplemental Figure 3 legend from previous page***

(**B**) Mean correlation coefficients (*r*), at different timepoints after injury, between transcriptomes of uninjured atlas RGCs and the injured RGCs (irrespective of cluster assignment), show that the changes in RGC transcriptomes over time after injury are non-linear. The difference between the time-points was significant by ANOVA (overall $F = 376.8$, $p < 0.0001$), and pairwise comparisons by posthoc LSD showed significant ($p < 0.001$) increase in correlation coefficient (*r*) at 1 week and 2 weeks, compared to the preceding 4 days time-point, after injury. Error bars = SEM.

(**C**) Percent of total injured RGCs unassigned by the iGraphBoost algorithm at each timepoint peaked at 4 and 7 days after injury, and then decreased by 2 weeks after injury.

(**D**) Mean correlation coefficients (*r*), at different timepoints after injury, between cluster assignment proportions of separate batches comprising each time-point, show that the highest variably between the batches was at 4 days after injury. Error bars = SEM.by 2 weeks after injury.

(**E-H**) RGC types segregate based on early and late onset of increased susceptibility to death after ONC, with some types that survived an early post-injury period (1st week) stabilizing by reducing rate of death (through 2nd week): Proportion by 1 week less than in atlas, and then by 2 weeks proportion less than at 1 week (early susceptible, non-stabilizing), includes 17 of the most susceptible (dark red-colored), along with medium-susceptible C5 and C37 (light red-colored) types in Fig. 3B (***E***); proportion by 1 week less than in atlas, and then by 2 weeks proportion higher than at 1 week (early susceptible, stabilizing), includes the medium-susceptible (light red-colored) types in Fig. 3B (***F***); proportion by 1 week higher than in atlas, and then by 2 weeks proportion less than in atlas (late susceptible, non-stabilizing), includes the medium-susceptible (light red-colored) types in Fig. 3B (***G***); and proportion by 2 weeks higher than in atlas (late susceptible, stabilizing), includes all the resilient (blue-colored) types in Fig. 3B (***H***). Days 2 and 4 time-points excluded from *E-H* due to higher batch variability per panel *D* above.

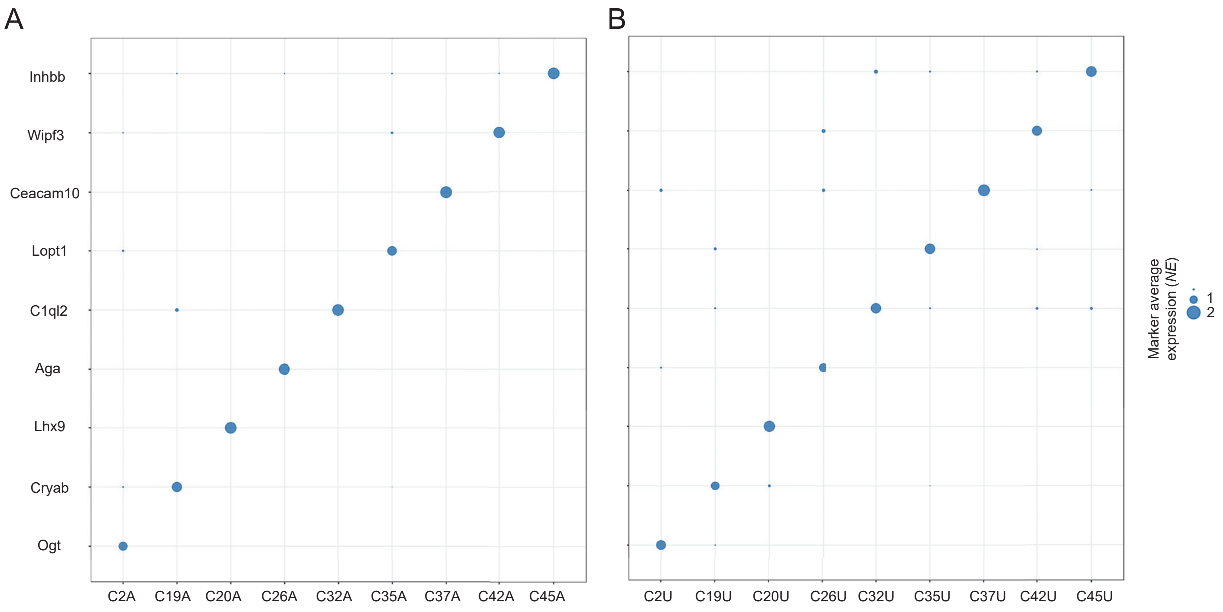


**Supplemental Figure 4. Originally unassigned injured RGCs express markers of the clusters to which they were assigned by the MapTo algorithm**

(**A-B**) Bubble plot of markers for the injured (2 weeks after ONC) RGCs that were assigned to the atlas RGC clusters by the iGraphBoost algorithm (***A***), and for the originally unassigned injured (2 weeks after ONC) RGCs that were assigned to the atlas RGC clusters in retrospect by the two-step MapTo algorithm (***B***). The same RGC type-specific markers are enriched in types assigned to the originally unassigned RGCs by the MapTo algorithm, as in the originally assigned RGC types. Because markers of the uninjured RGCs types changed expression in most of the RGC types by 2 weeks after injury (see Fig. 5B), markers that are specific to the injured types were used here. Bubble size scale bar indicates ***z***-scores of average normalized gene expression in a cluster relative to all other clusters. A = assigned originally by the iGraphBoost algorithm; U = unassigned originally but assigned in retrospect by the MapTo algorithm.

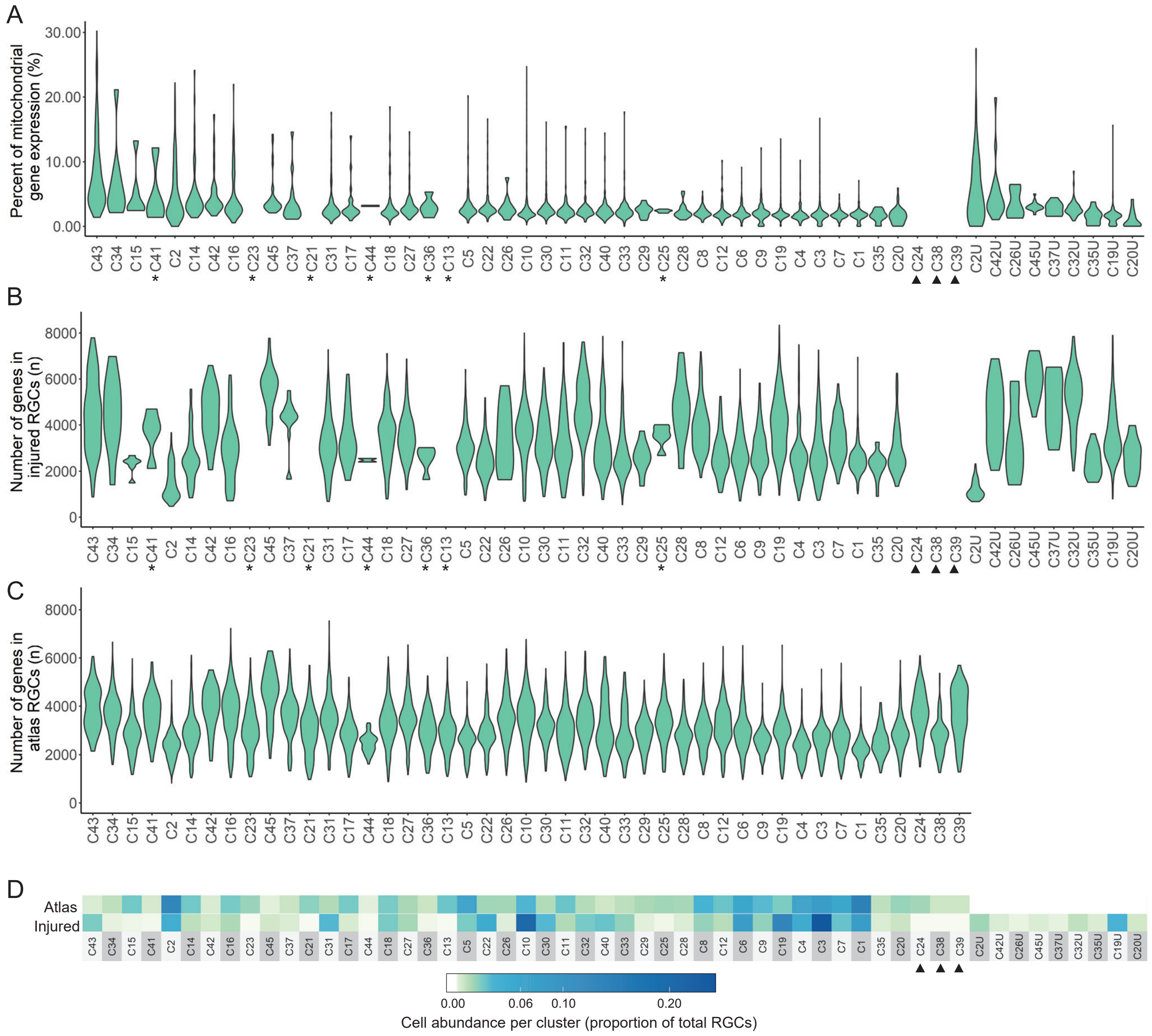


**Supplemental Figure 5. Global transcriptomic properties of the RGC types overrepresented in the originally unassigned injured RGCs are within the normal range compared to the other RGC types**

(**A**) Violin plots of the percentage of mitochondrial genes (which are indicative of cellular stress) per cluster of injured RGCs (at 2 weeks after ONC), showing that the percentage of mitochondrial genes of those RGC types which were overrepresented in the originally unassigned (U) injured RGCs (C2U, C19U, C20U, C26U, C32U, C35U, C37U, C42U, and C45U, shown at the right of the graph) are within the normal range compared to the percentage of mitochondrial genes in other RGC types. Cell types marked by an asterisk (*) represent those with 5 or less cells. Triangles mark those which did not survive 2 weeks after ONC, and thus contain 0 cells.

(**B**) Violin plots of the total number of genes per cluster (i.e., transcriptome size) of injured RGCs (at 2 weeks after ONC), showing that the transcriptome size of the RGC types which were overrepresented in the originally unassigned (U) injured RGCs (C2U, C19U, C20U, C26U, C32U, C35U, C37U, C42U, and C45U, shown at the right of the graph), are within the normal range compared to the other RGC types' transcriptome size.

(**C**) Violin plots of the total number of genes per cluster (i.e., transcriptome size) of uninjured atlas RGCs, for comparison to the changes in transcriptome size in RGC types after injury (shown in *B*).

(**D**) Abundance of uninjured atlas RGCs per cluster (upper row) and of injured RGCs per cluster (at 2 weeks after ONC; bottom row), showing that abundance of the RGC types which were overrepresented in the originally unassigned (U) injured RGCs (C2U, C19U, C20U, C26U, C32U, C35U, C37U, C42U, and C45U, shown at the right of the bottom row) is within the normal range, compared to the other RGC types' abundance.

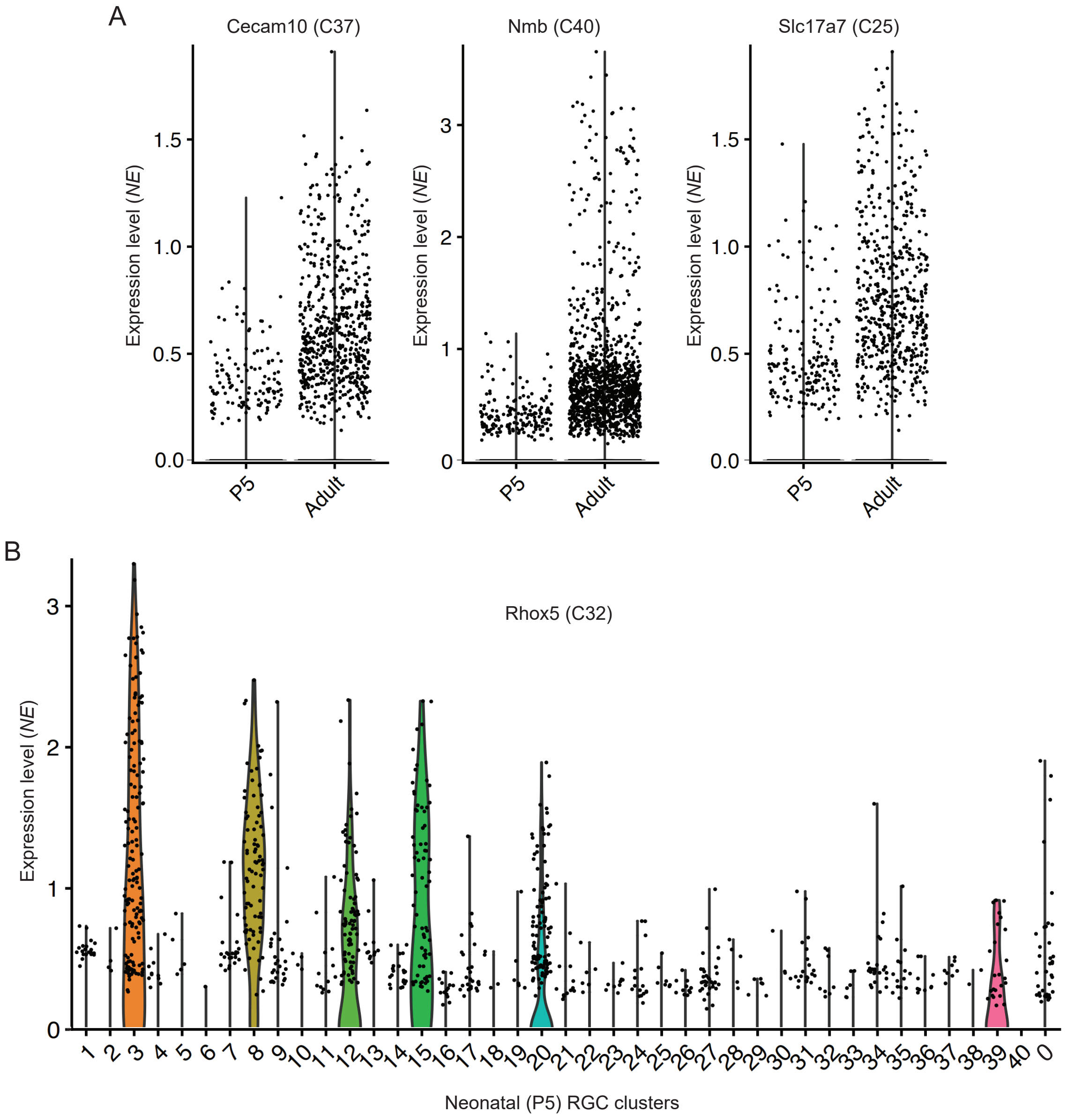


**Supplemental Figure 6. Cluster marker genes unshared between neonatal and adult RGC types are developmentally regulated**
(**A**) Violin plots showing that expression levels of Cecam10, Nmb, and Slc17a7 are all upregulated developmentally in the RGCs, from neonatal (postnatal day 5; P5) to adult timepoints, and thus did not qualify as cluster markers for the neonatal RGC types.
(**B**) Violin plots of Rhox5 expression in neonatal (P5) RGC clusters show enrichment in several clusters.

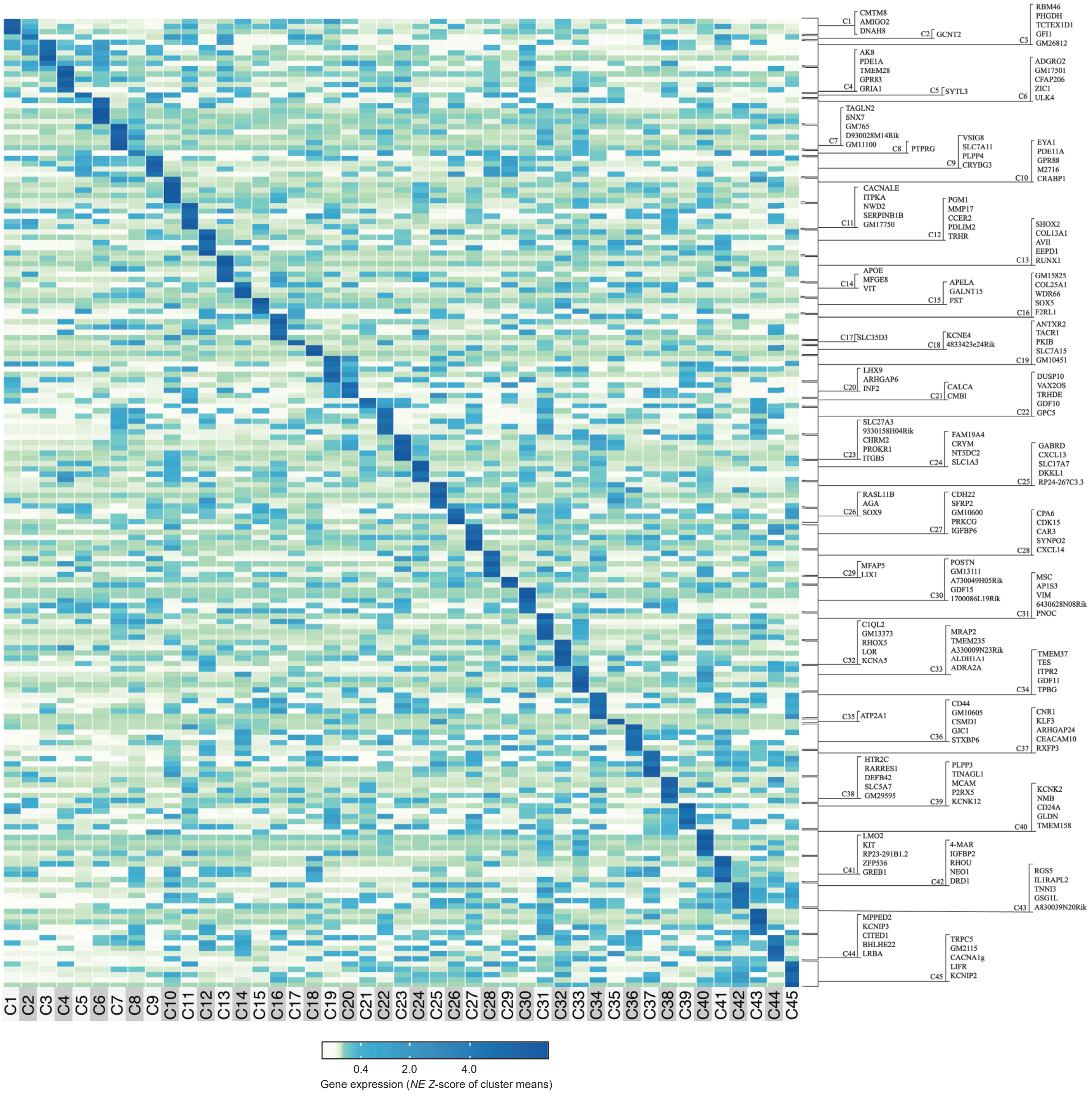


**Supplemental Figure 7. Heatmap signatures of cluster marker genes enriched in adult atlas RGC types**

Heatmap of adult atlas RGC clusters, with up to 5 enriched single gene cluster markers indicated on the right side. The enriched genes required the following selection criteria: Gene expression in a cluster $>0.05$ *NE* and $\geq 1.8$-fold relative to every other cluster, with enrichment *p*-value $\leq 0.05$ by both independent samples *t*-test (2-tailed) and the nonparametric independent samples Mann–Whitney *U* test. Color-coded scale bar of gene expression indicates *z*-scores after normalizing to all the clusters (see Methods). Other genes enriched in clusters that did not pass one or both statistical tests but met the first two criteria are listed in Supplemental Data 1.

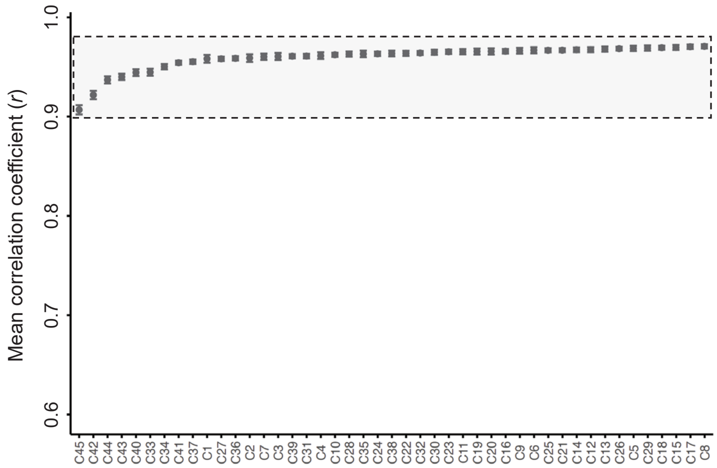


**Supplemental Figure 8. Downsampling atlas RGCs does not increase dissimilarity between the transcriptomes of atlas RGC types**

Mean Pearson's correlation coefficient (*r*) of the atlas RGC clusters compared to every other uninjured atlas RGC cluster, after atlas RGCs were downsampled to match the number of RGCs (8500) in the dataset from 2 weeks after ONC. Outlined in the shaded box *r* range here, is near-identical to the one shown in Figure 6A for the same analysis but using the full atlas RGC dataset (without downsampling).

| Gene | Adult Cluster | P5 Cluster |
|---|---|---|
| Apela | C15 | 3 |
| Pde1a | C4 | 7 |
| Prkcg | C27 | 16 |
| Postn | C30 | 18 |
| Fam19a4 | C24 | 29 |
| Mmp17 | C12 | 31 |
| Gpr88 | C10 | 32 |
| Zic1 | C6 | 34 |
| Prokr1 | C23 | 35 |
| Rhox5 | C32 | *N/A* |
| Ceacam10 | C37 | *N/A* |
| Nmb | C40 | *N/A* |
| Slc17a7 | C25 | *N/A* |

**Supplemental Table 1. Originally identified single gene RGC cluster markers**
Shows the single gene markers (13 in total) originally identified for adult RGC atlas clusters (by Tran et al. 2019)[4] and the clusters for which they were also identified as single gene markers in neonatal (postnatal day 5; P5) RGC atlas (by Rheaume et al., 2018)[5]. Three of the single gene markers, Ceacam10, Nmb, and Slc17a7, were developmentally upregulated and Rhox5 was enriched in several clusters (see **Supplemental Figure 6**), and thus they did not qualify as single gene markers for neonatal RGC clusters.

#### LEGENDS FOR SUPPLEMENTAL DATA

**Supplemental Data 1. Single gene cluster markers expression in adult RGC atlas clusters**
Genes uniquely enriched in adult RGC atlas clusters, and their normalized expression (*NE*), are shown. *NE* values are highlighted in beige in the columns that correspond to the clusters in which they are enriched. Up to 5 top gene markers per cluster with *p*-value ≤ 0.05 are specified in **Supplemental Figure 7**. Markers listed here also include genes that did not pass one or both of the statistical tests used (see Methods), but were still enriched ≥1.8-fold relative to every other cluster and had minimal expression ≥0.05 *NE*.

**Supplemental Data 2. R script of MapQuery implementation for comparative analysis of the integration mapping algorithms**
Downloadable and executable R script for implementation of the Seurat's MapQuery, in which the query dataset cells are mapped to the reference dataset cells for determining cluster assignment.

**Supplemental Data 3. Python script of scArches implementation for comparative analysis of the integration mapping algorithms**
Downloadable and executable Python script for implementation of scArches, in which the semi-supervised model scANVI in scArches is used to classify the query dataset cells based on the reference dataset cells for determining cluster assignment.

**Supplemental Data 4. R script of CIDER implementation for comparative analysis of the integration mapping algorithms**
Downloadable and executable R script for implementation of CIDER, in which the query dataset cells are integrated with the reference dataset cells by the assisted CIDER (asCIDER) clustering them together, and then percent of the query cells that were clustered correctly (i.e., within the clusters from which they originated prior to the query cells being set aside) is determined.

**Supplemental Data 5. R script of Symphony implementation for comparative analysis of the integration mapping algorithms**
Downloadable and executable R script for implementation of Symphony, in which the query dataset cells are mapped to the reference dataset cells for determining cluster assignment.
